# ScRNA-seq reveals tumor microenvironment remodeling induced by local intervention-based immunotherapy

**DOI:** 10.1101/2020.10.02.323006

**Authors:** Ashley R. Hoover, Kaili Liu, Christa I. DeVette, Jason R. Krawic, Connor L. West, Delaney Medcalf, Alana L. Welm, Xiao-Hong Sun, William H. Hildebrand, Wei R. Chen

**Affiliations:** Stephenson School of Biomedical Engineering, University of Oklahoma, Norman, OK, USA; Arthritis & Clinical Immunology Research Program, Oklahoma Medical Research Foundation, Oklahoma City, OK, USA; Department of Microbiology and Immunology, University of Oklahoma Health Sciences Center, Oklahoma City, OK, USA; Huntsman Cancer Institute, University of Utah, Salt Lake City, UT, USA

## Abstract

Laser immunotherapy (LIT) combines local photothermal therapy (PTT), to disrupt tumor homeostasis and release tumor antigens, and an intratumorally administered immunostimulant, N-dihydrogalactochitosan (GC), to induce antitumor immune responses. We performed single-cell RNA sequencing on tumor-infiltrating leukocytes of MMTV-PyMT mouse mammary tumors to characterize LIT-induced myeloid and lymphoid compartment remodeling. Analysis of 49,380 single cell transcriptomes from different treatment groups revealed that proinflammatory IFNα, IFNγ, and TNFα cytokine signaling pathways were enriched in both lymphoid and myeloid cells isolated from LIT-treated tumors. The CD4^+^ and CD8^+^ T cells in LIT treated tumors resided in an activated state while immune cells in untreated and PTT-treated tumors remained in a neutral/resting state. Additionally, monocytes recruited into the LIT-treated tumors were driven towards proinflammatory M1-like macrophage phenotypes or monocyte-derived dendritic cells. Our results reveal that LIT prompts immunological remodeling of the tumor microenvironment by initiating broad proinflammatory responses to drive antitumor immunity.

**STATEMENT OF SIGNIFICANCE:** Transcriptome profiling of tumor infiltrating leukocytes revealed that localized laser immunotherapy (LIT) greatly enhanced antitumor T cell activity by promoting proinflammatory myeloid cell responses within the tumor microenvironment. This manuscript demonstrates that LIT broadly stimulates antitumor immunity and has great potential to synergize with current immunotherapies to increase their efficacy.

## INTRODUCTION

A major challenge in cancer treatment is the failure of the host immune system to detect and destroy cancer cells (1), particularly metastasis, which causes 90% of cancer-related deaths. Effective therapies for metastatic cancers remain elusive (2). Cancer immunotherapy, which boosts the immune response to cancer, holds high promise for treating metastatic cancers (3). Various immunotherapies, such as cytokine and monoclonal antibody therapy (4,5) adoptive cell therapy (6), chimeric antigen receptor T-cell (CAR-T) therapy (7), tumor vaccines (8), and immune checkpoint therapy (ICT) (9), represent potent anti-cancer treatments. Unfortunately, most of these treatments rely on existing antitumor immune responses or prerequisite knowledge of target tumor antigens. Since most cancer patients have poor antitumor immunity, efficacy and response rates of current immunotherapies are low across various tumors in clinical studies (10,11).

A local intervention-based approach that induces systemic, long-term antitumor immunity represents an ideal therapy for metastatic cancers. Laser immunotherapy (LIT) was developed to achieve this objective. LIT has two components: 1) local photothermal therapy (PTT) for disrupting target tumor homeostasis and releasing tumor antigens (12,13), and 2) local administration of an immunostimulant, such as N-dihydrogalactochitosan (GC) (14,15). In preliminary clinical studies for patients with metastatic, treatment-recalcitrant breast cancer and melanoma, LIT successfully reduced and/or eliminated the treated primary tumors and untreated metastases in the lungs (16-18). Because of these promising outcomes in our pre-clinical and preliminary clinical studies, we are characterizing the underlying mechanism whereby LIT achieves powerful antitumor effects. Here, we hypothesized that LIT activates both innate and adaptive immune responses to foster sustained antitumor immunity.

To test this hypothesis, we evaluated the effects of LIT on breast tumors arising from the mouse mammary tumor virus-polyoma middle T (MMTV-PyMT) transgenic mice. MMTV-PyMT tumors have high penetrance and emulate the histological stages of human luminal type B breast cancer, providing an excellent immune competent model for studying the tumor microenvironment (TME) (19,20). Single-cell RNA sequencing (scRNA-seq) on human breast tumors revealed that tumor associated macrophages (TAMs), monocytes, and other myeloid and lymphoid cells are diverse in the TME (21). Therefore, we performed scRNA-seq on MMTV-PyMT tumors, to capture the complexity and heterogeneity of the TME following LIT treatment.

Using a combination of flow cytometry, T cell depletion, and scRNA-seq analysis, we revealed that LIT drives a global shift in immune cells towards various inflammatory states, fostering prolonged antitumor immunity. To evaluate LIT activation on the tumor-infiltrating leukocytes (TILs) in the TME, we analyzed scRNA-seq data on TILs from untreated (CTRL), PTT, GC, and LIT (PTT+GC) treated MMTV-PyMT tumors. Our results revealed that changes in the myeloid compartment potentiated adaptive immunity by increasing the activation of tumor-resident naïve, effector, and memory CD4^+^ and CD8^+^ T cells. LIT also induced a proinflammatory response in all the innate immune cells within the TME, increasing M1-like macrophages and prompting proinflammatory signals in M2-like macrophages. Together, our results demonstrate that LIT initiates broad changes within the TME, especially antitumor immune responses in both myeloid and lymphoid compartments, ultimately leading to systemic antitumor immunity.

## RESULTS

### LIT extends the survival of mice bearing MMTV-PyMT breast tumors

The therapeutic effects of LIT on breast tumors were determined using the MMTV-PyMT transgenic tumor model. Spontaneous MMTV-PyMT tumor cells were isolated from MMTV-PyMT transgenic mice as previously described (22) and were injected into the mammary fat pad of wild type female FVB mice (Figure 1A). When tumors reached a size of 0.5cm^3^, mice were divided into four treatment groups: Untreated controls (CTRL), PTT alone, GC alone, and LIT (PTT+GC). Treated animals were monitored up to 60 days following treatment, and euthanized once tumors reached ethical endpoints. LIT significantly reduced tumor growth and increased the survival rate compared to other groups (Figure 1B-C).

**Figure 1.**
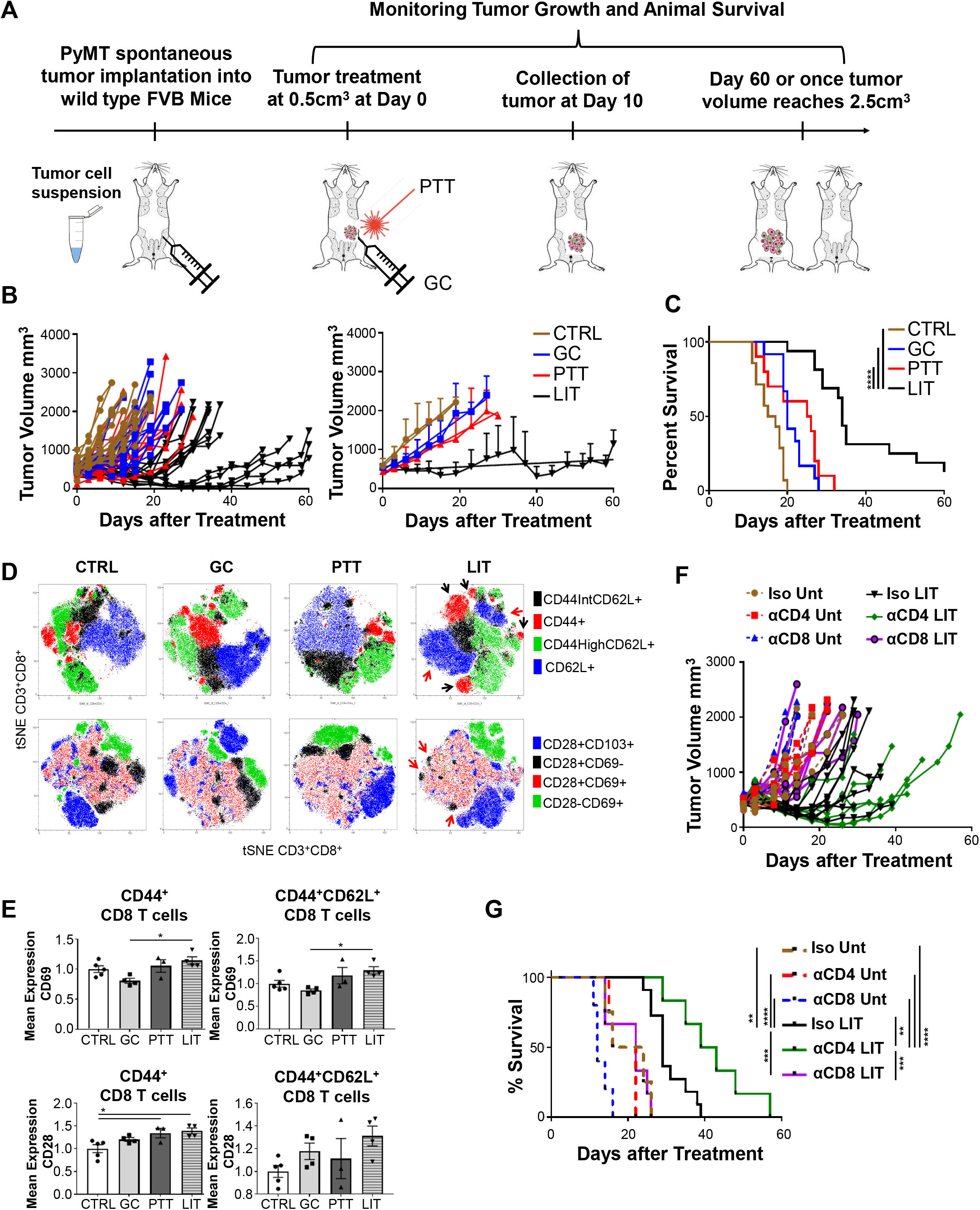
LIT treatment slows tumor growth by activating tumor-infiltrating CD8^+^ T cells. A: Schematic of MMTV-PyMT tumor implantation, treatment, and analysis. B: Individual and mean tumor size of mice after different treatments. C: Survival rates of tumor-bearing mice. Log-rank (Mantel-Cox) test was used for statistical analysis. D: T-SNE plots generated from 3 individual concatenated files in Flowjo of tumor-infiltrating CD8^+^ T cells from different treatment groups. Manual gates for CD44 and CD62L were overlayed onto the T-SNE plots. The colors indicate the expression of different cell surface marker combinations. T cells were analyzed 10 days after treatment. E: The MFI of CD69 and CD28 on CD8^+^ T cells, normalized to the CTRL, 10 days after treatment. One-way ANOVA was used for statistical analysis. F. Individual tumor size of untreated or LIT-treated mice that were injected with isotype antibodies or CD4^+^ or CD8^+^ depletion antibodies. G. Survival rates of untreated or LIT-treated tumor-bearing mice injected with isotype antibodies or depleted of CD4^+^ or CD8^+^ T cells. Log-rank (Mantel-Cox) test was used for statistical analysis.

Since cytotoxic T cells are crucial to antitumor immune responses, we evaluated the extent of cytotoxic T cell induction by LIT in the TME 10 days after treatment by flow cytometry. Thermal effects of laser irradiation reduce tumor size, resulting in significantly smaller tumors in the PTT and LIT groups than that in CTRL and GC groups (Figure S1A). For comparison, TIL cellularity was divided by tumor volume. The total normalized CD45^+^ cellularity/mm^3^ (Figure S1A), following the gating strategies shown in Figure S1B, showed no significant differences among different treatment groups. The effector cells (CD45^+^CD3^+^CD8^+^CD44^+^) and memory T cells (CD45^+^CD3^+^CD8^+^CD44^High^CD62L^+^) in LIT and GC groups had a trending increase, while the CD45^+^CD3^+^CD8^+^CD44^Low^CD62L^+^ and CD45^+^CD3^+^CD8^+^CD62L^+^ naïve T cells in LIT and GC groups were similar to that of CTRL and PTT groups (Figure S1A).

To further evaluate CD8^+^ T cell populations, using the gating strategy in Figure S1B, t-distributed Stochastic Neighbor Embedding (t-SNE) plots were generated from 3 concatenated samples from each group using FlowJo (Figure S1C). These plots revealed more diversity within the CD8^+^ T cells from LIT-treated tumors based on their distribution and number of distinct clusters (Figure 1D). For example, the CD62L^+^ cells in the LIT-treated tumors separated into two distinct groups (blue clusters, red arrows). The effector cells (red clusters, black arrows) from the LIT-treated tumors also separated into distinct clusters, with a much greater distance between them on the t-SNE plot, again suggesting more T cell diversity. There was also a large reduction in the proportion of CD28^+^CD69^-^ cells (Figure 1D, black clusters, red arrows) and an increase in the CD28^+^CD69^+^ cells in the LIT-treated tumors, suggesting the differentiation and activation of CD8^+^ T cells following LIT. The intensity and distribution of CD69^+^, CD28^+^, and CD103^+^ expression on CD8^+^ T cells were all increased in the LIT-treated tumors (Figure S1C). The mean expression levels of CD69, as determined by mean fluorescence intensity (MFI), on the CD3^+^CD8^+^CD44^+^ and CD3^+^CD8^+^CD44^+^CD62L^+^ (including both CD44^High^CD62L^+^ and CD44^Low^CD62L^+^ populations) T cell subsets, were statistically elevated in the LIT-treated tumors compared to GC alone, and trending up compared to CTRL and PTT (Figure 1E). CD28 was also significantly increased on the CD3^+^CD8^+^CD44^+^ cells and trending up on the CD3^+^CD8^+^CD44^+^CD62L^+^ T cell subsets (Figure 1E). These results indicate that the CD8^+^ T cells are activated in response to LIT, but this analysis did not fully explain the unique effects of LIT on tumor growth inhibition.

We next investigated the contributions of CD8^+^ and/or CD4^+^ T cells to the therapeutic effect of LIT, using depletion antibodies (Figure S1D), with >95% depletion confirmed by flow cytometry. Loss of CD8^+^ T cells after LIT treatment abrogated the therapeutic benefits of LIT, suggesting that cytotoxic T lymphocytes (CTLs) are critical for downstream responses following LIT (Figure 1F-G). Surprisingly, loss of CD4^+^ T cells following LIT treatment delayed tumor growth and prolonged survival in a manner superior to LIT alone (Figure 1G). These findings suggest that CD8^+^ T cells are imperative to mounting an anti-tumor response, but CD4^+^ T cells may serve as an avenue for immune inhibition post-LIT exposure in this model. We therefore proceeded with scRNA-seq to provide a transcriptomic explanation for the immunological changes driven by LIT.

### ScRNA-seq reveals diversity of tumor-infiltrating leukocytes induced by LIT

To investigate a broad spectrum of immune cell alterations in the TME, scRNA-seq was performed, following the workflow depicted in Figure S1E. Mice bearing MMTV-PyMT tumors were divided into four treatment groups (n=4 per group). Nine days after treatment, tumor-infiltrating leukocytes (TILs) were pooled from mice in each group and enriched. First, all 49,380 TILs (11,584 CTRL; 14,071 PTT; 12,501 GC; 11,224 LIT) were integrated for scRNA-seq. Unsupervised clustering analysis was performed using Seurat R package (23). Using the default shared-nearest neighbor (SNN) resolution (0.5), t-SNE and Uniform Manifold Approximation and Projection (UMAP) plots unveiled 20 (0-19) distinct cell clusters, representative of both myeloid and lymphoid cell lineages (Figure 2A-B). The unsupervised analysis of the top 10 differentially expressed genes (DEGs) from each cell confirmed that the TILs were successfully delineated into 20 distinct clusters as visualized by the heatmap (Figure 2C).

**Figure 2.**
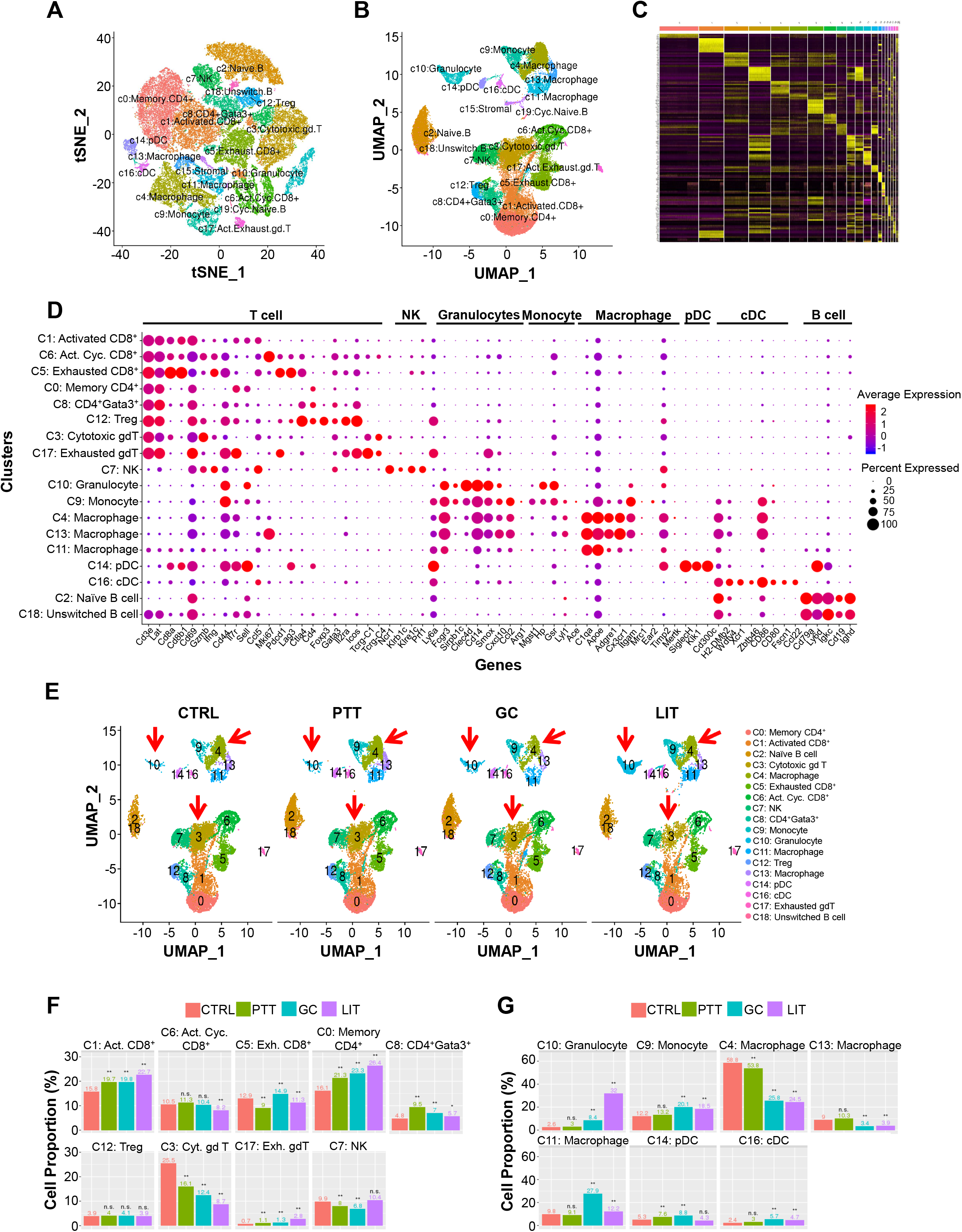
ScRNA-seq of CTRL, GC, PTT, and LIT treated tumors creates a tumor-infiltrating immune cell atlas. A: Two-dimensional visualization of single cell clusters using method of t-distributed Stochastic Neighbor Embedding (t-SNE) from integrated immune cells of different treatment groups. B: Two-dimensional visualization of single cell clusters using method of Uniform Manifold Approximation and Projection (UMAP) from integrated immune cells of different treatment groups. C: Heatmap of gene expression for the top 10 DEGs in each cluster (yellow bars). D: Validation of unsupervised clustering using traditional immune cell genes. E: UMAP plots of the immune cell atlas from individual treatment groups. F. Proportion of lymphoid cells in different clusters compared to total lymphocytes in different treatment groups. Statistical analysis was performed using proportion test function (prop.test) in R. G: Proportion of myeloid cells in different clusters compared to total myeloid cells in individual treatment groups. Statistical analysis was performed using proportion test function (prop.test) in R.

To accurately annotate the immune cell clusters, we used a combination of the cluster identity predictor (CIPR) (https://aekiz.shinyapps.io/CIPR/) which is linked to the Immunological Genome Project (ImmGen) and a publicly available cell specific gene list (24). With the top 5 scores generated by CIPR, we identified the following TIL populations: T cells, γδ-T cells, Tregs, B cells, pre-B cells, granulocytes, monocytes, macrophages (Mφ), dendritic cells (DCs), and natural killer (NK) cells (Figure S2A). Stromal cell and B cell clusters (15 and 19) were excluded from the rest of the analysis. With CIPR, cells in cluster 6 cannot be clearly distinguished as either T or B cells by the top 5 scores. To overcome this limitation, we relied on a series of commonly used immune cell genes to distinguish and confirm the cell type in each cluster (Figure 2D). The dot plots show the average expression levels and percentages of specific genes commonly used to identify immune cell subtypes (Figure 2D). The specificity of these selected genes is illustrated with the correlation plot for the entire cell-derived gene expression patterns (Figure S2B), indicating that their expressions correlate with and represent specific cell clusters.

By combining the gene expression and cluster analysis we annotated all cell clusters (Figure 2A-D). For example, cells in cluster 6 were activated, proliferating CD8^+^ T cells, with high expression of *Cd8a, Cd8b1, Gzmb, IFNg*, and cell cycling marker *Mki67*. The combination of unsupervised clustering using high throughput signature annotation and common traditional immune cell genes for validation were mutually supportive and complementary (Figures 2C-D, Figures S2A-B).

Separating the UMAP of integrated cells (Figure 2B) into four individual plots (Figure 2E), we discovered that several clusters were significantly different in the LIT-treated tumors compared to that of CTRL and PTT groups (Figure 2E, red arrows). We also calculated the immune cell proportions in individual compartments (Figure 2F-G) and the proportions of immune cells in individual clusters against total TILs (Figure S2C). Specifically, LIT increased the proportions of activated CD8^+^ T cells (cluster 1) and memory CD4^+^ T cells (cluster 0), and decreased cytotoxic γδ-T cells (cluster 3) (Figure 2F). As shown in Figure 2G, LIT increased the proportions of granulocytes (cluster 10) and decreased Mφ (cluster 4). These findings support our hypothesis that LIT induces broad changes to TILs in the TME.

### LIT alters lymphoid cell composition and gene expression in the TME

We next explored transcriptomic changes within specific T cell subsets, since diversity of T cell subtypes in the TME have profound effects on antitumor immunity. Our scRNA-seq data demonstrated that activated CD8^+^ T cells and memory CD4^+^ T cells were significantly increased in LIT-treated tumors (Figure 2F). To further analyze these cell types, we performed higher resolution re-clustering of the lymphoid cells, excluding B cells, and obtained 21 clusters (clusters 0-20), as shown in Figure 3A. Expressions of common lymphoid cell markers, including *Cd3e, Cd4, Cd8b1, Foxp3, Mki67*, and *Ncr1*, were examined and mapped to cell clusters using UMAP plots (Figure S3A), and annotation was performed (Figure S3B). The increased resolution revealed the subtypes of lymphoid cells that experienced significant changes (Figure S3C, red arrows) with the activated CD8^+^ T cells from cluster 1 in Figure 2F becoming clusters 0 and 3 in Figure 3B. LIT significantly upregulated naïve CD8^+^ T cells (cluster 0) and central-memory CD4^+^ T cells (cluster 9), as shown in Figure 3B. Clusters 5 and 6 both contained naïve CD4^+^ T cells, with cluster 5 resembling stimulated T cells and cluster 6 unstimulated (Figure 3B). Additionally, LIT decreased the cell ratio of cluster 1 (cytotoxic γδ-T cells) and induced a small proportion of IL-17^+^IFNγ^+^ CD8^+^ T cells (cluster 19; Figure 3B). The appearance of naïve T cells suggests that LIT can actively recruit these cells to the TME and potentially generate *de novo* T cell responses, not just enhance existing T cells responses. This is significant as LIT has the potential to generate better, more diverse T cell pool that limits exhaustion and immune evasion.

**Figure 3.**
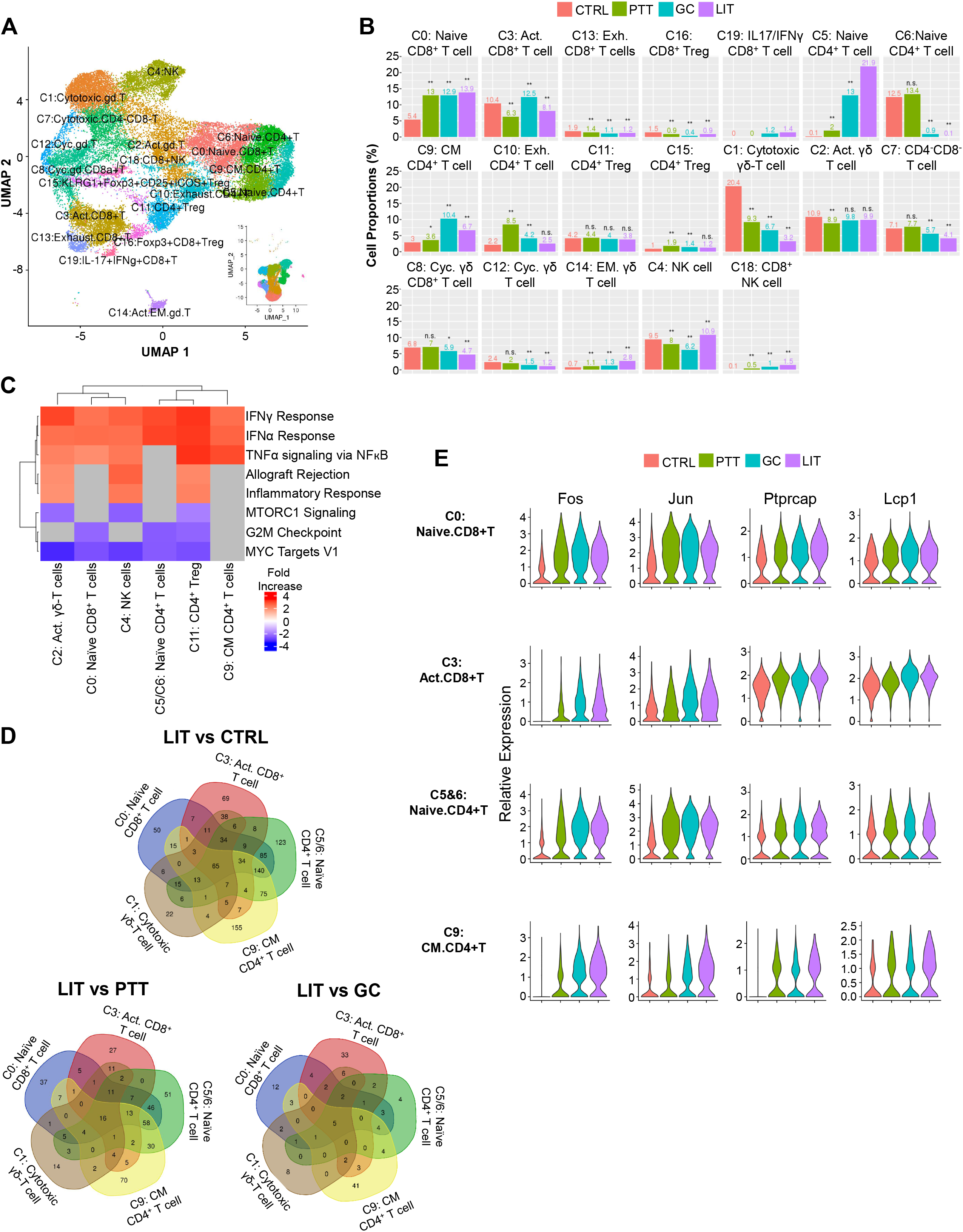
LIT treatment enriches proinflammatory signaling pathways in TILs. A: UMAP of re-clustered lymphoid cells with an increased SNN resolution (from 0.5 to 0.7). The insert is the clusters of lymphoid cells with the lower SNN (0.5) in Figure 2B. B: Proportions of re-clustered lymphoid cells in different treatment groups. C: Heatmap of pathway enrichment for lymphoid cell clusters induced by LIT using GSEA. D: Venn diagrams showing the number of DEGs and commonly expressed genes in clusters 0, 3, 5/6, 1, and 9. E: Expression levels of 4 representative DEGs identified in the Venn diagrams from cluster 0, 3, 5/6, and 9.

To determine how the lymphoid cells are activated, we performed gene set enrichment analysis (GSEA). Using differentially expressed genes (DEGs), we focused on the comparison between LIT and CTRL groups. As shown in Figures 3C and S3D-E, LIT significantly upregulated IFNα and IFNγ signaling pathways of activated γδ-T cells (cluster 2), naïve CD8^+^ T cells (cluster 0), NK cells (cluster 4), the combined naïve CD4^+^ T cells (cluster 5 and 6), CD4^+^ Tregs (cluster 11), and central memory CD4^+^ T cells (cluster 9). mTORC1, G2M checkpoint, and Myc targets pathways were downregulated by LIT in these cell clusters (Figure 3C and S3D-E). The signaling pathway analysis demonstrates that LIT promotes the antitumor type I/II IFNs and TNF pathways across different lymphoid cell types.

To find the common DEGs in five separate lymphoid cell clusters from different comparison groups, we generated Venn diagrams (Figure 3D). In these selected lymphoid subtypes, 65, 16, and 5 commonly shared DEGs were found in LIT vs CTRL, LIT vs PTT, and LIT vs GC, respectively (Figure 3D, Table S1). We also generated violin plots for a series of conserved DEGs for LIT vs CTRL in these selected T cell types and found that LIT significantly promoted the expression of several genes involved in T cell activation, function, and survival (Figure 3E and S3F and Table S1). For example, LIT promoted the expression of Gimap proteins, *Dusp1 (MKP-1), Isg15*, and *Ptprc*, which are involved in T cell survival, activation/function, and death (Figure S3F) (25-28). LIT also induced the expression of *Fos, Jun, Ptprcap*, and *Lcp1* (Figure 3E), which are associated with T cell receptor (TCR) signaling (29-31), indicating that T cells in LIT-treated tumors are responding to their cognate antigens. LIT also initiated changes in several other genes with less known functions in T cells, as listed in Table S1. In summary, our data reveal that LIT induces TCR signaling within T cells.

### LIT promotes naïve and memory T cell activation in the TME

To confirm that T cells in LIT-treated tumors were activated, we performed cell trajectory inference (CTI) analysis, using R package monocle2 (32), on CD8^+^ (Figure 4A-D, S4A) and CD4^+^ T cells (Figure 4E-H and S4D). The CD8^+^ T cells were divided into 9 states (Figure S4A) with the naïve CD8^+^ T cells distributed on the right branches, including states 4, 5, and 6 (Figure 4A and S4A). Non-naïve subtypes of CD8^+^ T cells were distributed on the left branches, including states 1, 2, 3, 7, 8, and 9 (Figure 4A and S4A). When separated based on different treatments, the cell trajectory patterns were shifted by GC and LIT from upper states (1, 2, 3, 5, 9) to lower states (6, 7, 8), as shown in Figure 4B. To determine if the shift was indeed due to T cell activation, we generated volcano plots of the top DEGs of CD8^+^ T cells between LIT-treated and CTRL tumors (Figure S4B). The genes highly regulated in CD8^+^ T cells from LIT-treated tumors were associated with T cell activation (Figure S4B). Thus, we infer that cells in the upper branches are in a resting/neutral state (blue circle) and the cells in lower branches are in an activated state (red circles), as shown in Figure 4C-D. By examining individual CD8^+^ T cell clusters, we observed a shift in the CD8^+^ T cells in clusters 0, 3, 13, and 16 towards the activated states in GC and LIT-treated tumors (Figure 4D). Overall, the proportion of CD8^+^ T cells in the activated state increased dramatically in the LIT-treated tumors (Figure S4C).

**Figure 4.**
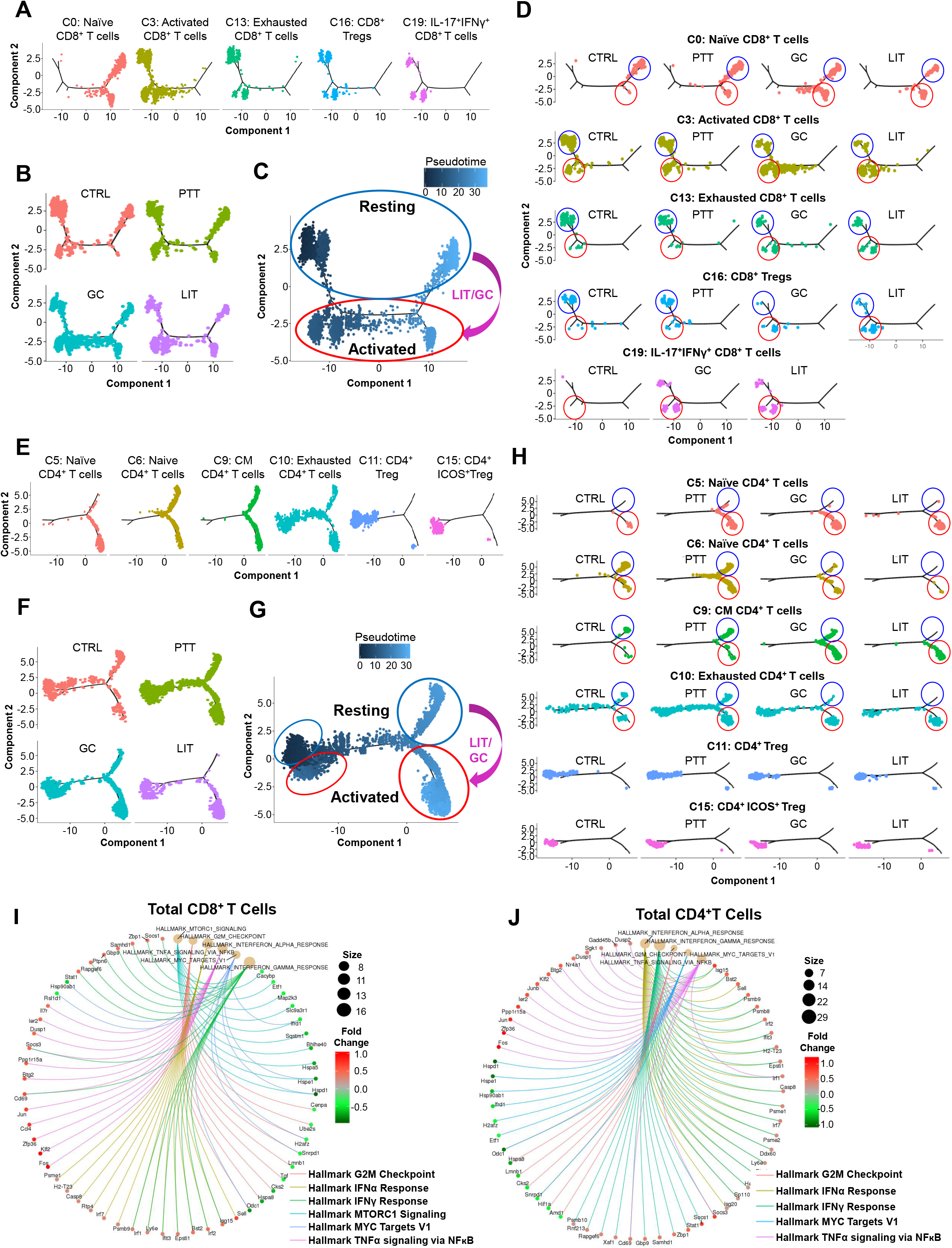
T cells from LIT-treated tumors reside in activated states. A: Branched trajectory of CD8^+^ T cells in different clusters separated according to each annotated cell type. B: Branched trajectory of CD8^+^ T cells separated according to treatment group. C: LIT induced transitioning (purple arrow) pattern for CD8^+^ T cells from a resting state (blue circle) to an activated state (red circle). D: Branched trajectories of CD8^+^ T cell subtypes in different treatment groups. Cells in blue circles are in a resting state and cells in red circles are in an activated state. E: Branched trajectory of CD4^+^ T cells in different clusters separated according to each annotated cell type. F: Branched trajectory of CD4^+^ T cells separated according to treatment group. G: LIT/GC induced transitioning (purple arrow) pattern for CD4^+^ T cells from a resting state (blue circle) to an activated state (red circle). H: Branched trajectories of CD4^+^ T cell subtypes in different treatment groups. Cells blue circle are in a resting state and cells in red circles are in an activated state. I: Pathway enrichment for all CD8^+^ T cell clusters induced by LIT using GSEA. J: Pathway enrichment for all CD4^+^ T cell clusters induced by LIT using GSEA.

We also performed CTI analysis on CD4^+^ T cells (Figure 4E and S4D). CD4^+^ T cells were divided into 11 states (Figure S4D). Naïve and central memory CD4^+^ T cells were distributed on the right branches, which included states 6 through 11 (Figure 4E and S4D). Exhausted CD4^+^ T cells showed ubiquitous expression throughout the trajectory. The Tregs were localized at the left branches, in an opposite pattern to that of naïve CD4^+^ T cells (Figure 4E). When separated based on treatment groups, the cell trajectory patterns were shifted by GC and LIT from the upper states to lower states (Figure 4F), similar to that of CD8^+^ T cells (Figure 4B). To confirm this switch was also due to T cell activation, volcano plots from DEGs of CD4^+^ T cells between LIT-treated and CTRL tumors were generated (Figure S4E). The genes associated with T cell activation were upregulated in LIT-treated tumors (Figure S4E). Based on the gene expression data, we labeled the cells residing in the lower states (red circle) as activated and cells residing in the upper states (blue circles) as resting/neutral (Figure 4G-H). The proportion of cells residing in the resting/neutral state versus the activated state was significantly different in the GC and LIT-treated tumors compared to that in the CTRL and PTT-treated tumors (Figure S4F).

GC and LIT shifted CD4^+^ T cells towards the activated state in clusters 5, 6, 9, and 10 (Figure 4H). Using our re-clustering analysis of the lymphoid cells, we noted that even though clusters 5 and 6 contained both naïve CD4^+^ T cells, the cells in cluster 5 were activated while cells in cluster 6 were resting/neutral. This likely explains the dramatic shift in these two populations by LIT (Figure 3B) as nearly all the T cells in cluster 5 reside in the activated state along with very few T cells from cluster 6 in the LIT-treated tumors (Figure 4H). Interestingly, only in the LIT-treated tumors we observed that all the central memory and exhausted T cells (cluster 9) resided in the activated state (Figure 4H).

To determine how LIT is globally effecting CD8^+^ and CD4^+^ T cell activation, we performed GSEA on total CD8^+^ or CD4^+^ T cells from LIT-treated and CTRL tumors. LIT significantly upregulated proinflammatory signaling pathways IFNα, IFNγ, and TNFα and downregulated Myc targets and G2M checkpoint regulators in both CD8^+^ (Figure 4I) and CD4^+^ (Figure 4J) T cells.

Thus far our results demonstrate that LIT significantly enriches T cell activation in the TME. However, GC alone does not directly activate T cells (data not shown). To uncover how changes within the innate immune system allow for global T cell activation, we further analyzed modifications of the myeloid cells following LIT.

### LIT induces a proinflammatory phenotype in myeloid cells

Within the immunosuppressive TME of an established tumor, TAMs generally resemble M2-like Mφ and inhibit migration and function of T cells (33,34). TAMs play an essential role in the progression and metastasis of breast cancer (35,36). With the dynamic role of TAMs, it is not surprising that TAMs residing in the TME have diverse functions (21,37). To explore the heterogeneity of the myeloid cells, we used a higher SNN resolution (0.7) to re-cluster these immune cells. Using principal component analysis (PCA), we obtained 19 clusters (clusters 0-18), annotated using a series of commonly used immune cell genes (Figure 5A and S5A). We filtered out 4 non-myeloid clusters and kept 15 myeloid clusters for further analysis (Figure 5A). Comparing UMAPs of different treatment groups, we observed notable differences in the composition of granulocytic myeloid derived suppressor cells (G-MDSCs), Mφ, and DCs in the LIT-treated tumors (Figure S5B, red arrows). The proportions of immune cells in the 15 clusters were calculated for each group (Figure 5B). LIT enhanced the proportions of G-MDSC (clusters 3 and 8) and M1-like Mφ (cluster 7), but significantly decreased M2-like Mφ (clusters 0 and 1), as shown in Figure 5B. Phenotypically, M1-like Mφ and M2-like Mφ may be distinguished by cell surface expression of *Itgam* and *Adgre1* (Figure S5C).

**Figure 5.**
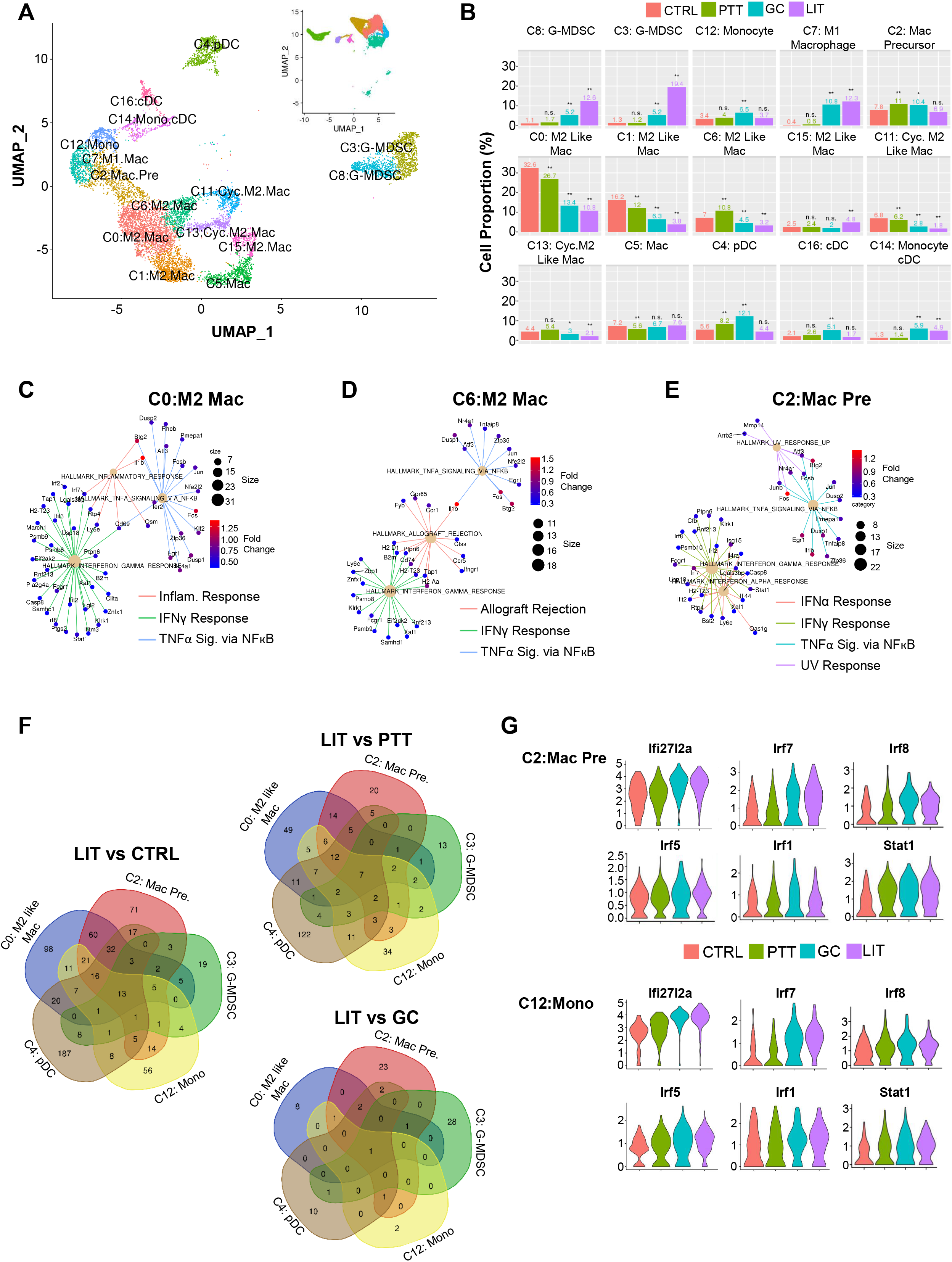
LIT primes tumor-resident innate immune cells to proinflammatory states. A: UMAP of re-clustered myeloid cells with an increased SNN resolution (from 0.5 to 0.7). The insert is the clusters of myeloid cells with the lower SNN (0.5) in Figure 2B. B: The proportions of re-clustered myeloid cells from individual treatment groups. Statistical analysis was performed using proportion test function (prop.test) in R. C-E: Pathway enrichment for representative myeloid cells (clusters 0, 6, and 2) induced by LIT using GSEA. F: Venn diagrams showing the number of DEGs and commonly expressed genes identified in each myeloid cell cluster. G: Expression levels of select genes upregulated in cluster 2 and 12.

In addition to the myeloid cell proportions, we also investigated the alterations in their activation states. For the Mφ clusters, we calculated the DEGs and performed GSEA. First, we compared Mφ in LIT-treated and CTRL tumors. LIT increased proinflammatory responses of Mφ in clusters 0, 6, and 2, compared to that in untreated tumors (Figure 5C-E). Specifically, these pathways include IFNγ, TNFα, and inflammatory responses of M2-like Mφ in cluster 0 (Figure 5C), and IFNγ, TNFα, and allograft rejection pathways of M2-like Mφ in cluster 6 (Figure 5D). In cluster 2, Mφ precursors, IFNα, IFNγ, TNFα, and UV response pathways were upregulated by LIT (Figure 5E). Next, we combined all M2-like Mφ clusters for GSEA and discovered LIT significantly upregulated IFNα and IFNγ signaling pathways, and decreased hypoxia, TNFα, and UV response pathways (Figure S5D). These results demonstrate that even though LIT-treated tumors contain M2-like Mφ, they are responding to type I IFNs and exhibiting a more proinflammatory phenotype than M2-like Mφ from CTRL tumors.

Next, we analyzed G-MDSC gene expression in LIT treated tumors using gene ontology (GO) analysis. DEGs comparison between LIT and other groups was not practical due to low cell numbers in CTRL group (Figure 5B). GO analysis of G-MDSCs in LIT-treated tumors revealed 6 highly enriched pathways involved in neutrophil degranulation and interferon signaling (Figure S5E). These cytokine pathways are associated with proinflammatory responses and can enhance immunity. Additionally, signaling pathways of IL-10 and IL-13, the cytokines traditionally associated with MDSC immunosuppression, were also enriched in the G-MDSCs (Figure S5E). Interestingly, when we compared G-MDSCs in cluster 3 to cluster 8 in LIT-treated tumors, LIT enriched IFNα response genes in cluster 3, but reduced the TNFα, UV response, apoptosis, hypoxia, epithelial to mesenchymal transition, p53, mTORC1, and MYC target pathways (Figure S5F). This is of particular interest because mTOR, hypoxia, and glycolysis signaling pathways are interconnected and they are key to MDSC suppressive function (38,39). The reduction in these signaling pathways suggests a diminished immunosuppressive function of G-MDSCs in cluster 3 compared to that in cluster 8. Despite the increase in G-MDSCs, other key cell types (T cells, DCs, and TAMs) are activated and functional in the TME. Our findings suggest that the immune-suppressive function of G-MDSCs within the LIT-treated tumors is reduced.

Finally, we evaluated genes that did not comply to specific pathway networks with Venn diagrams of common DEGs across clusters 0, 2, 12, 3, and 4 in different groups: LIT vs CTRL, LIT vs PTT, and LIT vs GC. We found 13 conserved DEGs in LIT vs CTRL, 7 in LIT vs PTT, and 1 in LIT vs GC (Figure 5F, Table S2). Violin plots of these genes demonstrated that LIT significantly increased the expression of *Ifi27l2a* and *Irf7* (Figure 5G). LIT also increased the expression of *il1b, Btg2, Fos, Junb, Zfp36, Trim30a, Oasl2, Slfn5*, and *Sp100*, and inhibited expression of *Ninj1* (Figure S6A). Particularly interesting is the high expression of *Irf7* in monocytes (cluster 12) from GC- and LIT-treated tumors (Figure 5G), since *Irf7* drives monocyte differentiation into Mφ (40). Cells in cluster 2, the Mφ precursors, showed significantly upregulated *Stat1* and displayed trending increases in the proportion of cells expressing *Irf1, Irf8*, and *Irf5*, genes expressed in activated M1-like Mφ (Figure 5G), likely explaining why LIT-treated samples contain more M1-like Mφ (40,41). These results imply that LIT not only changes the behavior of existing M2-like Mφ, but also recruits and differentiates monocytes to become proinflammatory cells, enhancing antitumor immunity.

### LIT shifts the TME towards a proinflammatory phenotype, promoting monocyte differentiation into M1-like macrophages

To determine the effect of LIT on the TME in driving monocytes towards M1-like Mφ and/or shifting M2-like Mφ to M1-like, we conducted cell CTI analysis and discovered 7 major states. M2-like Mφ clusters were shown to reside within states 1, 2 and 7 (Figures 6A and S6B). M2-like Mφ in clusters 0, 1, and Mφ precursors in cluster 2 (Figure 6A), show a broader trajectory covering states 1, 2, 4, 6, and 7 (Figure S6B-C), indicating that these Mφ were in a more plastic state and could be driven towards an M1- or M2-like phenotype depending on the environmental signals they receive. Since the trajectories look identical between all four treatment groups, we conclude that cluster 0 and 1 are not the predominant source of M1-like Mφ (Figure S6C). However, these Mφ clusters could be targets for M2 to M1 conversion therapeutics in combination with LIT or other immunotherapies.

**Figure 6.**
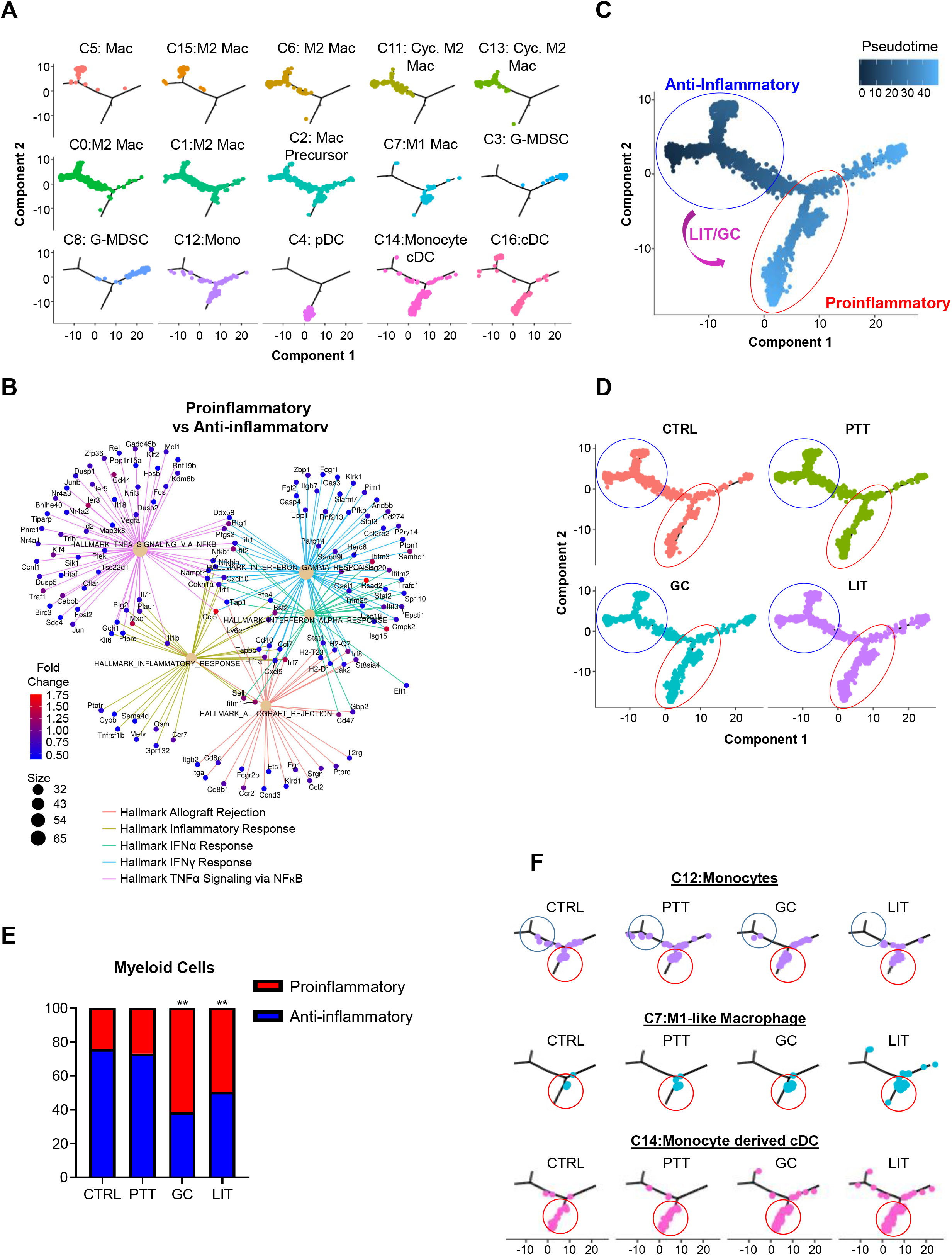
LIT promotes tumor-infiltrating monocytes differentiation to become proinflammatory cell types. A: Single cell trajectory inference of re-clustered myeloid cells in different treatment groups. B: Pathway enrichment for DEGs by comparing proinflammatory trajectories to anti-inflammatory using GSEA. Clusters 7, 14, and 16 (proinflammatory) were compared to clusters 5, 6, 11, 13, and 15 (anti-inflammatory). C: LIT/GC induced transitioning (purple arrow) pattern for myeloid cells from an anti-inflammatory state (blue circle) to a proinflammatory state (red circle). D: Branched trajectories of selected myeloid cells in different treatment groups. Blue circles indicate anti-inflammatory (pro-tumor) trajectories while red circles indicate proinflammatory (anti-tumor) trajectories E: Cell proportions from anti-inflammatory and proinflammatory myeloid trajectories in different treatment groups. Statistical analysis was performed using proportion test function (prop.test) in R. ** means *p* < 0.01. F: Branched trajectories of selected myeloid cells (M1 macrophages of cluster 7, Monocytes of cluster 12, and Monocyte-derived DCs of cluster 14) in different treatment groups.

The DCs (clusters 4, 14, and 16), G-MDSCs (clusters 3 and 8), and M1-like Mφ (cluster 7) were shown to reside in states 3, 4, 5, and 6 (Figure 6A and S6B). We performed GSEA to compare the states of DCs and M1-like Mφ to M2-like Mφ (clusters 5, 15, 6, 11 and 13), M1-like Mφ to M2-like Mφ, and lastly G-MDSCs to M2-like Mφ. Although clusters 0 and 1 were also annotated as M2-like Mφ, they were not chosen due to their ubiquitous distributions along the trajectories. This comparison revealed significant enrichment of IFNα, IFNγ, TNFα, and inflammatory responses in DCs and M1-like Mφ (Figure 6B and S6D). Interestingly, as shown in Figure S6E, G-MDSCs (clusters 3 and 8) displayed a significant upregulation of IFNα, IFNγ, and TNFα inflammatory responses, as well as immunosuppressive responses (apoptosis, hypoxia, and IL6/Jak/Stat3 signaling) compared to M2-like Mφ. To make LIT treatment more successful in eliminating breast tumors, it is of future interest to combine this treatment with a G-MDSC inhibitor and/or anti-PD-L1 checkpoint therapy.

Based on the GSEA, we designated trajectory branches at states 1, 2, and 7 as anti-inflammatory, and states 4, 5, and 6 as proinflammatory (Figure 6C). Interestingly, when we separated the trajectory in Figure 6C into 4 treatment groups, we noticed that GC and LIT-treated tumors had a significantly larger proportion of cells residing in the proinflammatory state than in the anti-inflammatory state (Figure 6D-E). This demonstrates that LIT drives myeloid cells towards a proinflammatory phenotype.

To determine if the overall proinflammatory TME in GC and LIT-treated tumors contributes to the differentiation of the recruited monocytes towards M1-like Mφ and/or monocyte-derived DCs (moDCs), we compared the cell trajectories of monocytes, M1-like Mφ, and moDCs among the four treatment groups. Although LIT did not change the monocyte cell proportion (cluster 12 in Figure 5B), the monocyte cellular distribution was changed along the trajectory. The cells in the upper-left anti-inflammatory sides (states 1, 7, and 2) shifted towards the downward/lower-right proinflammatory sides (states 4, 5 and 6), in which the M1-like macrophages and moDCs reside (Figures 6F and S6B) suggesting that monocytes are the predominant source of M1-like Mφ.

By using both Seurat and monocle2 analysis procedures, we conclude that LIT actively remodels the TME by shifting the cell composition and gene signatures from anti-inflammatory to proinflammatory. Additionally, we believe that the predominant source of the M1-like Mφ is monocytes, not a result of M2 to M1 Mφ conversion. Lastly, the proinflammatory response generated in the myeloid compartment allows for the activation of T cells in the TME to potentiate antitumor immunity.

## DISCUSSION

In this study, scRNA-seq provided unbiased interrogation of immune cells residing within the TME of the MMTV-PyMT tumor model, to elucidate the immunological changes initiated by LIT from a cellular and transcriptional vantage point. We analyzed 49,380 TILs and characterized the cell types, cell states, and dynamic cellular transitions. LIT significantly changed the landscape of multiple immune cell types of both myeloid and lymphoid cells by increasing or decreasing several key immune cell populations, upregulating immune activation genes, and inducing a type I/II IFN cytokine signature in the TME. While GC also induced similar immune responses, GC alone, similar to PTT alone, could not slow or stop tumor growth. This indicates that increasing the inflammatory capacity of immune cells alone is not enough, and that it must be combined with disrupted tumor homeostasis to initiate antitumor functions.

Initial analysis of T cell activation and T cell depletion revealed that, in the MMTV-PyMT breast tumor model, CD8^+^ T cells are essential for the therapeutic effect of LIT (Figures 1F-G). This may not be the case for every tumor model, and our investigations on the necessity of CD4^+^ T cells following LIT in other tumor models are ongoing. Further transcriptome analysis using scRNA-seq revealed that effector and memory T cells from LIT-treated tumors were responding to type I and II IFNs and transitioned to an activated state based on gene expression and CTI analysis (Figure 3E and Figures 4A-H). Interestingly, LIT dramatically increased the numbers of naïve CD4^+^ and CD8^+^ T cells in the TME, suggesting that LIT triggers active recruitment of naïve T cells (Figure 3B). That implies that LIT turns the TME into a tertiary lymphoid organ (TLO), which is a lymphoid-like structure where active antigen presentation and subsequent B/T cell activation occur (42). This is of critical importance because TLOs are associated with higher TIL numbers, increased responses to ICT, and positive outcomes of cancer patients (42,43). Supporting this notion, the naïve T cells residing in the TME following LIT are responding to antigens as they have increased expression of genes involved in TCR signaling and activation, and have transitioned into an activated state (Figure 3E and Figure 4). Furthermore, TCR signaling and activation in naïve T cells portents that LIT promotes the generation of *de novo* antitumor T cell responses, not just enhances the existing anti-tumor T cell responses. This is important as immunologically “cold” tumors with low numbers of T cells could become “hot” following LIT, increasing the numbers of tumor specific T cells. Lastly, increasing the numbers and activity of tumor-infiltrating T cells makes LIT an encouraging treatment option to synergize with ICT, which relies on the number of T cells in the TME (44) and the activation state of these T cells (45). We speculate that the increased quality and improved quantity of T cells induced by LIT will synergize with ICT, allowing for reduced ICT doses, minimizing the potential for autoimmune side effects, and broadly increasing ICT efficacy across different cancers. Thus far, we have not noticed any autoimmune sequelae after LIT treatment, an observation that makes LIT with adjuvant ICT therapy particularly attractive.

A majority of the TAMs residing in the TME resemble M2-like Mφ and exhibit immunosuppressive functions (46). Our data revealed that in CTRL tumors the vast majority of myeloid cells are M2-like Mφ, with smaller populations of DCs and G-MDSCs (Figure 5B). LIT-treated tumors exhibited a dramatic increase in M1-like Mφ, and conversely a decrease in the two most prominent subtypes of M2-like Mφ (Figure 5B) that exhibit a broad trajectory by CTI analysis (Figure S6C). Although M2-like Mφ from clusters 0 and 1 hovered between both M1- and M2-like states, they do not appear to be the predominant source of M1-like Mφ as all the treatment groups have cellular distribution that looks identical (Figure S6C). However, it is likely these Mφ retain plasticity and could be influenced by the local TME to become either M1- or M2-like Mφ. Since clusters 0 and 1 are the predominant M2-like Mφ populations within the TME, they make promising candidates for M2 to M1 conversion therapeutics (47), which could increase the clinical efficacy of LIT in breast cancer patients with high M2-like Mφ infiltration.

We next analyzed monocyte and Mφ precursors in the TME to determine if these cells give rise to a majority of the M1-like Mφ. Gene expression and CTI analyses revealed that the monocytes recruited into the TME after LIT were poised to become Mφ and moDCs (Figure 5B and 6F). Supporting this notion, monocytes from the LIT-treated tumors have increased IRF7 expression (Figure 5G), which drives Mφ differentiation (40). In turn, the Mφ precursors (cluster 2) in LIT-treated tumors were poised to become M1-like Mφ based on their increased expression of *Stat1, Irf1, Irf5, Irf7, and Irf8* (Figure 5G), genes enriched in M1 Mφ (41). CTI analysis also revealed that LIT significantly increased the numbers of myeloid cells residing in the proinflammatory state and decreased the numbers of cells residing in the anti-inflammatory state (Figure 6C-E).

We also observed a significant increase in granulocytic cells which were identified as G-MDSC (Figure 5B). Increased number of G-MDSC is another indicator that LIT drives a proinflammatory response, as MDSCs are normally recruited to high inflammation areas to promote immune suppression and wound healing. MDSCs are known immune suppressors and contribute to the establishment of the primary TME and a metastatic niche. Notably, G-MDSCs are less immunosuppressive than their monocytic-MDSC counterparts and are more similar to neutrophils (48). Gene ontology analysis revealed that the G-MDSCs from LIT-treated tumors responded to type I/II IFNs and also expressed the classic immunosuppressive signatures such as IL-10 and IL-13 signaling pathways (Figure S5E). Despite this immunosuppressive gene signature, LIT-treated tumors were enriched with activated CD4^+^ and CD8^+^ T cells (Figures 3 and 4), suggesting that the immunosuppressive function of the G-MDSCs may be reduced following LIT. However, we cannot discount the immunosuppressive ability of G-MDSCs as these cells express high levels of PD-L1, which may explain why a single application of LIT is not curative in this model (Figure 1B-C). Therefore, LIT combined with G-MDSC inhibitors, or anti-PDL1 ICT, may greatly improve the efficacy of LIT in highly immunosuppressive tumors.

While the current study provided unequivocal support for the systemic antitumor effects of LIT, it also revealed some *de novo* mechanisms to be explored in order to further improve the efficacy of LIT. For example, ICT is a promising candidate to be synergized with T cells activated by LIT. Depending on the tumor type, M2-like Mφ and MDSC inhibitors make promising combinations to enhance the curative potential of LIT in highly immunosuppressive TMEs such as in triple negative breast cancer. By directly disrupting primary tumor homeostasis, we reversed the pro-tumor TME. Its ability to be customized with other appropriate therapies, dependent on the tumor immune profile, could make LIT an effective personalized immunotherapy for metastatic cancer patients.

## ACKNOWLEDGMENTS

This work was supported in part by the National Cancer Institute (R01CA205348 to WRC and 1R01CA223245 to ALW) and the Oklahoma Center for the Advancement of Science and Technology (HR16-085). We would like to thank the Flow Cytometry Core Facility and Genomics Core Facility at the Oklahoma Medical Research Foundation for providing flow cytometry services and assisting in scRNA-seq analysis. The authors (WRC, ARH, KL) also appreciate the generous support of Dr. and Mrs. Lee and Sherry Beasley, Dr. and Mrs. J. Michael and Kathy Steffen, and Dr. and Mrs. Damon and Heather Johnston. We thank Samuel Kelting for computational assistance. We also thank Dr. Samuel Siu Kit Lam from Immunophotonics for scientific input.

## AUTHOR CONTRIBUTIONS

WRC, WHH, ARH conceived the project

ARH, CID, KL, JRK, CLW, DM performed the experiments

ARH, KL, CID analyzed the data

ARH, KL, WRC completed the original draft of the manuscript

WHH, CID, XHS, ALW, JRK reviewed and edited the manuscript

ALW provided tumor samples

## KEY RESOURCES TABLE

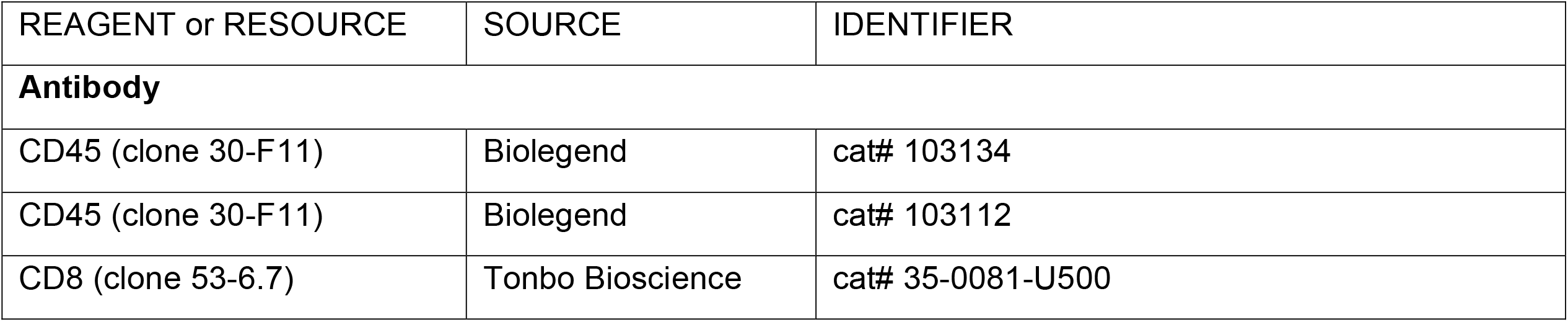
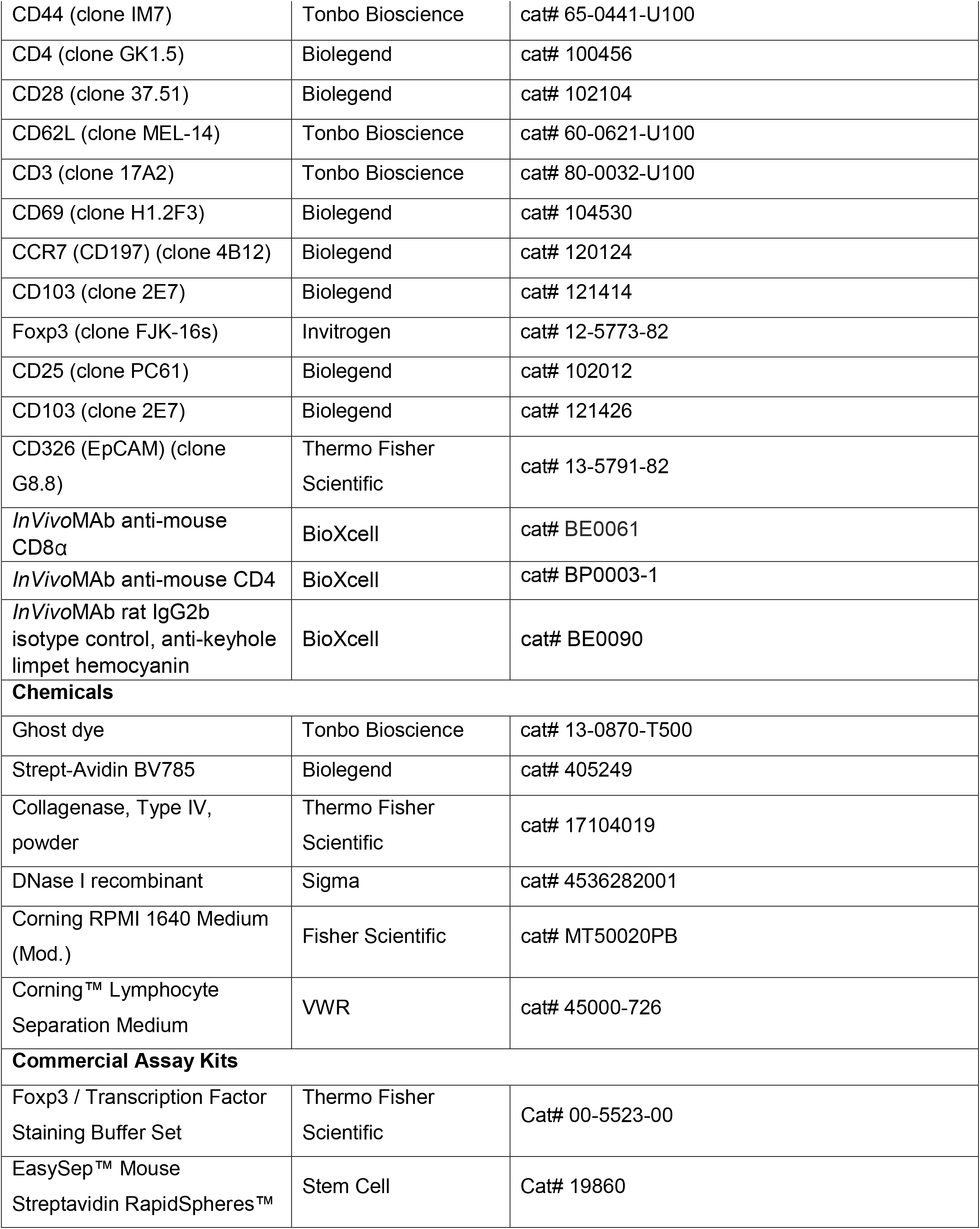
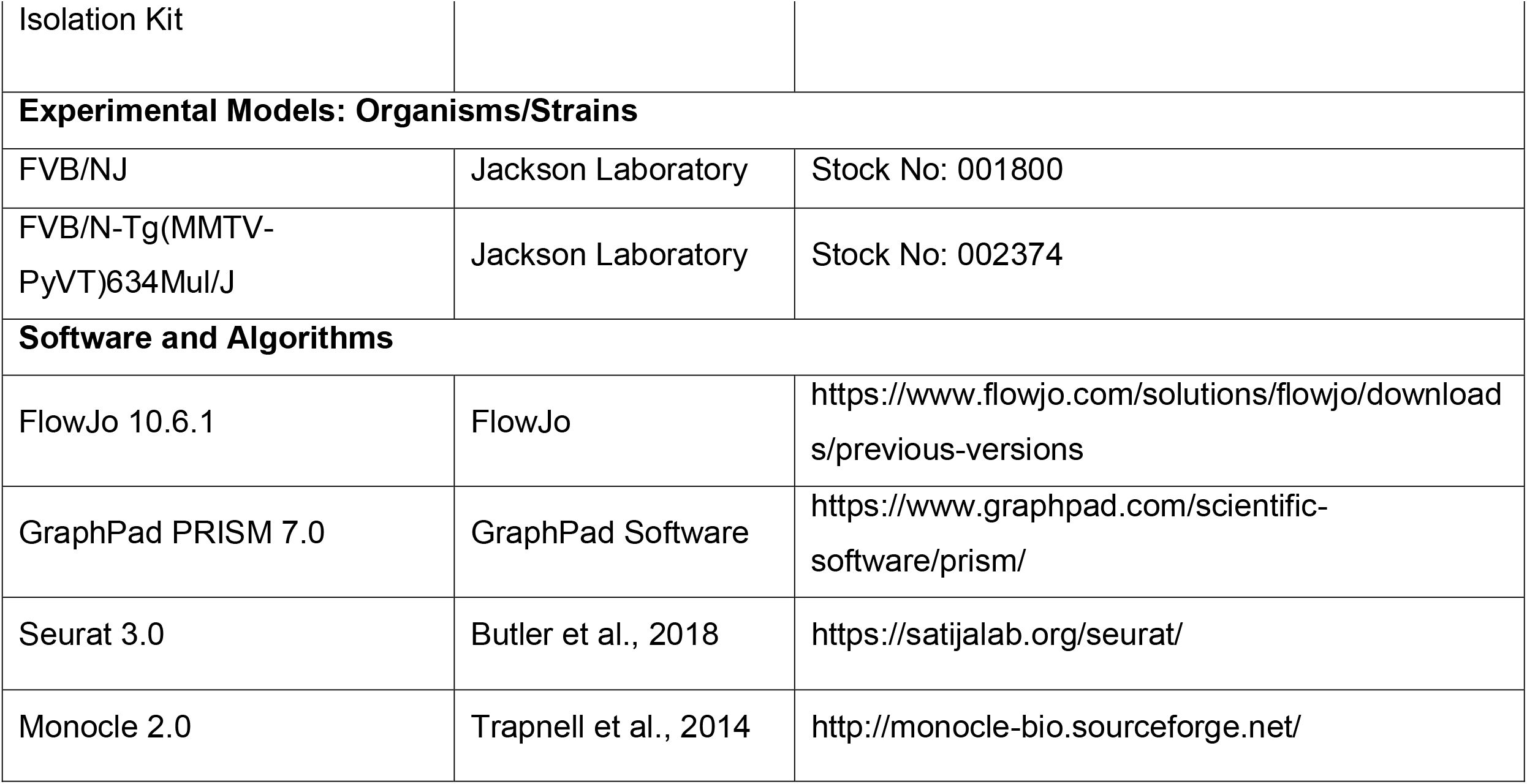

## CONTACT FOR REAGENTS AND RESOURCE SHARING

Further information and requests for resources and reagents should be directed to and will be fulfilled by the lead contact, Wei R. Chen.

## STUDY APPROVAL

Experimental procedures and handling of mice were performed in accordance with the University of Oklahoma Health Sciences Center (OUHSC) and Oklahoma Medical Research Foundation (OMRF) Institutional Animal Care and Use Committee (IACUC) regulations and approved protocols.

## METHODS

### MICE

Female FVB/NJ wild-type (stock #001800) and FVB/N transgenic MMTV-PyMT (stock #002374) mice were purchased from Jackson Laboratories and housed according to institutional guidelines at the University of Oklahoma Health Sciences Center. All studies were approved and adhered to OUHSC Institutional Animal Care and Use Committee protocols (#17-032-HCL).

### SYNGENEIC TUMOR CELL TRANSPLANTATION

PyMT murine breast tumor organoids were isolated from FVB/ N-Tg(MMTV-PyVT)634Mul/J mice as previously described (22). Briefly, cells were incubated overnight in mammary epithelial cell media (DMEM/F12 supplemented with 10% fetal bovine serum, 100 U/mL penicillin-streptomycin, 5ug/mL insulin-transferrin-selenium, 1ug/mL hydrocortisone, 10ng/mL mouse epidermal growth factor (EGF), and 50 ug/mL gentamicin (Sigma-Aldrich, St. Louis, MO)). Cells were washed twice with Hepes Buffered Saline (HBSS), trypsinized, and resuspended to a concentration of 1×10^5^ cell per 20uL. Cells were injected into the mammary fat pad of wild type FVB mice without clearing. The incision was closed with Vetbond tissue adhesive (3 M).

### TREATMENT OF MOUSE TUMORS

When tumors reached 0.5 cm^3^, the tumor site was shaved and mice received one of four treatments: CTRL (without treatment), GC alone, PTT alone, and LIT (PTT+GC). The mice in GC group received an intratumoral injection of 1% GC in 0.1 ml solution (Immunophotonics, Inc.) For mice in PTT group, the tumor was treated by an 805-nm laser (AngioDynamics, Latham, NY) with a power density of 1 W/cm^2^ for 10 minutes, using an optical fiber with a diffusion lens (Pioneer Optics, Bloomfield, CT) to delivery uniform light distribution on the treatment surface. For the mice in LIT group, the tumor was treated by the laser (1 W/cm^2^ for 10 minutes), followed by an intratumoral injection of GC (0.1 ml at 1%). Ten days after treatment, tumors were collected from selected mice in each group. The remaining mice were observed for up to 60 days or when the tumors reached a size of 2.5 cm^3^, or reached ethical endpoints.

### IN VIVO DEPLETION OF T CELLS

Once tumors reached 0.4cm^3^, 400μg of anti-CD4, -CD8, or isotype antibody (BioXcell) was injected i.p. Twenty-four hours after antibody injection, tumors were treated with LIT or left untreated. Three days after LIT treatment, mice were injected with 200μg antibody, i.p. and maintained on this dose biweekly until tumors reached a size of 2.5cm^3^ or reached ethical endpoints.

### TUMOR HARVEST

Tumor tissues were isolated, minced, and digested with Collagenase IV and DNase I at 37°C for 20-30 minutes in RPMI. After enzymatic digestion, immune cells were enriched using lymphocyte separation medium (Corning). The enriched cells were then used for flow cytometry analysis.

### FACS AND CONVENTIONAL FLOW CYTOMETRY

All samples were run on the LSR II and data was analyzed using FlowJo v10.6.1 (BD Biosciences, San Jose, CA). The cells used for scRNA-seq were sorted using the MoFlo sorting system. The antibodies used for flow cytometry are listed in the key resources.

### SINGLE-CELL RNA SEQUENCING

#### Sample Preparation

In each of the four treatment groups, immune cells from four mice were pooled together for single cell sequencing. Tumor tissues were isolated, minced with scalpels, and they digested with Collagenase IV and Dnase I at 37°C for 20-30 minutes. After enzymatic digestion, immune cells were enriched using lymphocyte separation medium. The enriched cells were then subjected to magnetic bead separation to remove the EpCAM^+^ cells. The EpCAM-depleted cells were stained with antibodies against CD45 and a viability dye. Live CD45+ cells were sorted using MoFlo and then processed for scRNA-seq via a 10× Genomics platform according to the manufacturer’s instructions (10× Genomics). Paired-end RNA-seq was performed via an Illumina NovaSeq 6000 sequencing system. Sequencing reads were processed using the 10× Genomics CellRanger pipeline and further analyzed with the Seurat R package. The effect of mitochondrial gene representation and the variance of unique molecular identifier (UMI) counts were regressed out from the data set prior to analysis. Gene expression signatures defining cell clusters were analyzed after aggregating samples from all 4 treatment groups (CTRL, PTT, GC, LIT). The raw data from scRNA-seq experiments in this manuscript can be found in the NCBI’s Gene Expression Omnibus database (GSE150675).

#### ScRNA-seq Library Generation

Droplet-based 3’ end scRNA-seq was performed by encapsulating sorted live CD45^+^ tumor-infiltrating immune cells into droplets using the 10x Genomics platform. cDNA libraries were prepared using Chromium Single Cell 3’ Reagent Kits v3 according to the manufacturer’s protocol (10x Genomics). The generated scRNA-seq libraries were sequenced using an Illumina NovaSeq 6000.

#### Alignment, Barcode Assignment, UMI Counting

The Cell Ranger (https://support.10xgenomics.com/single-cell-gene-expression/software/pipelines/latest/what-is-cell-ranger) was used to conduct sample demultiplexing, barcode processing, and single-cell 3’ counting. The Linux command *cellranger mkfastq* was applied to demultiplex raw base call (BCL) files from the illumine NovaSeq6000 sequencer, into sample-specific fastq files. Then, fastq files for each sample were processed with *cellranger count*, which was used to align samples to mm10 genome, filter and quantify.

#### Data Preprocessing with Seurat R Package

Seurat-guided analyses (https://satijalab.org/seurat/vignettes.html) were used to preprocess and integrate datasets from different treatment groups (Butler et al., 2018). Genes that were expressed in less than 5 cells or cells that expressed less than 8000 and more than 6000 genes were excluded. Also, cells that expressed less than 512 and more than 92600 counts or with a mitochondria percentage over 10% were excluded. Most variable genes were identified using the FindVariableFeatures function by setting feature numbers as 2000. Principal component analysis (PCA) was performed on the scaled matrix (with most variable genes only) using the first 30 principal components (PCs). Both tSNE and UMAP dimensional reductions were carried out using the first 20 PCs to obtain two-dimensional representations of the cell states. For clustering, we used the function FindClusters that implements a shared nearest neighbor (SNN) modularity optimization-based clustering algorithm on 30 PCs with resolution 0.5 for default analysis. For individual myeloid and lymphoid cell cluster analysis, 40 PCs and resolution 0.7 were used to gain more granularity and separate more clusters for immune cell subtypes.

#### Identification of Cluster-specific Genes and Marker-based Classification

Cells clusters were obtained by using the FindClusters function of the Seurat R package, which identifies clusters through an SNN modularity optimization–based algorithm. The biological cell type identities of each cluster were annotated with the assistance of an immune-cell scoring algorithm comparing the differentially expressed gene (DEG) signatures obtained from Seurat with the Immunological Genome Project Database (ImmGen) (24). This *in silico* cell type annotation was verified by using known canonical immune cell genes.

#### Individual Myeloid and Lymphoid Population Analysis

To increase the granularity, we manually used 40 PCs and 0.7 of SNN to separate myeloid and lymphoid clusters. This higher resolution approach allowed us to differentiate subsets of granulocytes, macrophages, dendritic cells, and T cell clusters.

### SINGLE CELL TRAJECTORY ANALYSIS

Monocle2 was used to construct the single cell trajectory landscape of tumor immune infiltrates including myeloid cells, CD8^+^ T and CD4^+^ T cells (32). Seurat object was imported into Monocle2. Although monocle2 can provide cell type identification, we used Seurat to obtain cell type annotation information and generate trajectory using the *differentialGeneTest* function with the argument *fullModelFormulaStr* setting Seurat derived clusters. The data dimension reduction for trajectory was performed using the DDRTree algorithm in the Monocle package. Further information about Monocle can be found in Monocle’s manual and vignette on Bioconductor.

### GSEA

Differential Gene Expression was obtained by using the FindMarkers function in Seurat. MAST was used as the methodology. Ranked gene lists were then analyzed for gene set enrichment by using the clusterProfiler R package (49). Hallmark gene sets (H) from the Molecular Signature Database (MSigDB, https://www.gsea-msigdb.org/gsea/msigdb) were used in these analyses (50). Curated gene sets (C2) and immunologic gene sets (C7) were also examined but not shown.

### STATISTICAL ANALYSES

Evaluations for tumor growth and FACS data were conducted using one-way Anova. MAST test was used for analyzing differential gene expression in selected cell clusters. P values less than or equal to 0.05 were considered statistically significant throughout (*, p ≤ 0.05; **, p ≤ 0.01).

## DATA AND SOFTWARE AVAILABILITY

The accession number for the scRNAseq data reported in this paper is GEO: GSE150675 (https://www.ncbi.nlm.nih.gov/geo/query/acc.cgi?acc=GSE150675). Analysis of such data can also be available upon request.

