## Supplementary material for "ScRNA-seq reveals tumor microenvironment remodeling induced by local intervention-based immunotherapy": Suppl. Figure Legends

**Supplemental Figure Legends**

**Figure S1.** **T cell activation by LIT (related to Figure 1)**

A: Tumor size, total TILs, and CD8^+^ T cell cellularity divided by tumor volume.

B: Gating strategy for CD4^+^ and CD8^+^ T cells.

C: T-SNE plots generated from 3 individual concatenated files in Flowjo of tumor-infiltrating CD8^+^CD3^+^ T cells. T-SNE plots are representative heat maps of the intensity and distribution of CD28, CD103, and CD69 within the CD8^+^CD3^+^ T cells.

D: Schematic of depletion of CD4^+^ and CD8^+^ T cells and LIT treatment of tumor-bearing mice.

E: Flowchart for TIL isolation and scRNA-seq after different treatments.

**Figure S2.** **Annotation of tumor-infiltrating immune cells using scRNA-seq data (related to Figure 2).**

A: Cell type annotation by CIPR database.

B: Gene-gene correlation heatmap of selected immune cell genes.

C: Proportion of lymphocytes and myeloid cells compared to total TILs. Statistical analysis was performed using proportion test function (prop.test) in R.

**Figure S3.** **Analysis of lymphoid cells using scRNA-seq after treatment (related to** **Figure 3)**.

A: An increased SNN resolution (from 0.5 to 0.7) UMAP of re-clustered lymphoid cells overlaid with common lymphoid cell gene expression, excluding B cells.

B: Validation of unsupervised re-clustering using traditional lymphoid cell genes.

C: UMAP of re-clustered lymphoid cells with an increased SNN resolution (from 0.5 to 0.7) for different treatment groups. Red arrows indicate cell clusters with significant change induced by LIT when compared to CTRL.

D-E: Pathway enrichment for representative lymphoid cell clusters induced by LIT using GSEA. Gene sets were annotated for the pathways using Hallmark gene sets of Molecular Signatures Database (MSigDB).

F. Expression levels of 7 representative DEGs identified in the Venn diagrams from cluster 0, 3, 5/6, and 9.

**Figure S4.** **T cells from LIT-treated tumors are in an activated state (related to** **Figure 4)**.

A: Branched trajectory of CD8^+^ T cells separated according to each state, generated using monocle2.

B: Volcano plots of differential gene expression of CD8^+^ T cells (LIT versus CTRL) in clusters 0, 3, and 13.

C: CD8^+^ T cell proportions from resting and activated trajectories in different treatment groups. Statistical analysis was performed using proportion test function (prop.test) in R. ** means *p* < 0.01.

D: Branched trajectory of CD4^+^ T cells separated according to each state. States were generated using monocle2.

E: Volcano plots of differential gene expression of CD4^+^ T cells (LIT versus CTRL) in clusters 5/6, 9, and 10.

F: CD4^+^ T cell proportions from resting and activated trajectories in different treatment groups. Statistical analysis was performed using proportion test function (prop.test) in R. ** means *p* < 0.01.

**Figure S5.** **Tumor resident innate immune cells primed to become proinflammatory by LIT (related to Figure 5)**.

A: Validation of unsupervised re-clustering using traditional myeloid cell genes.

B: UMAP of re-clustered myeloid cells separated by each treatment group: CTRL, PTT, GC and LIT. Red arrows indicate cell clusters with significant change induced by LIT when compared to CTRL.

C: Delineation of M1- vs M2-like macrophages by expression levels of *Itgam* and *Adgre1* in myeloid cell clusters.

D: Pathway enrichment for remaining M2-like macrophage clusters (not included in clusters 0 and 6) induced by LIT using GSEA.

E: Gene ontology for G-MDSCs in LIT-treated tumors using the Reactome pathway database.

F: Pathway enrichment for LIT-induced G-MDSCs in clusters 3 versus cluster 8 using MSigDB.

**Figure S6.** **Inducing tumor-infiltrating monocytes to become proinflammatory cell types by LIT (related to** **Figure 6)**.

A: Expression levels of selected common LIT-regulated genes in selected myeloid cell clusters.

B: Branched trajectories of all myeloid cells separated by each state, generated using monocle2.

C: Branched trajectories of M2 macrophages of clusters 0 and 1, macrophage precursor cells of cluster 2 in different treatment groups.

D: Pathway enrichment for DEGs by comparing G-MDSCs with M2-like macrophages using MSigDB.

E: Pathway enrichment for DEGs by comparing M1-like macrophages with M2-like macrophages using MSigDB.

**Supplemental Tables**

**Table S1:** Differentially expressed genes regulated by LIT in selected lymphoid subtypes.

**Table S2:** Differentially expressed genes regulated by LIT in selected myeloid cell subtypes.
