## Supplementary material for "ScRNA-seq reveals tumor microenvironment remodeling induced by local intervention-based immunotherapy": Suppl. Figures

### Slide 1
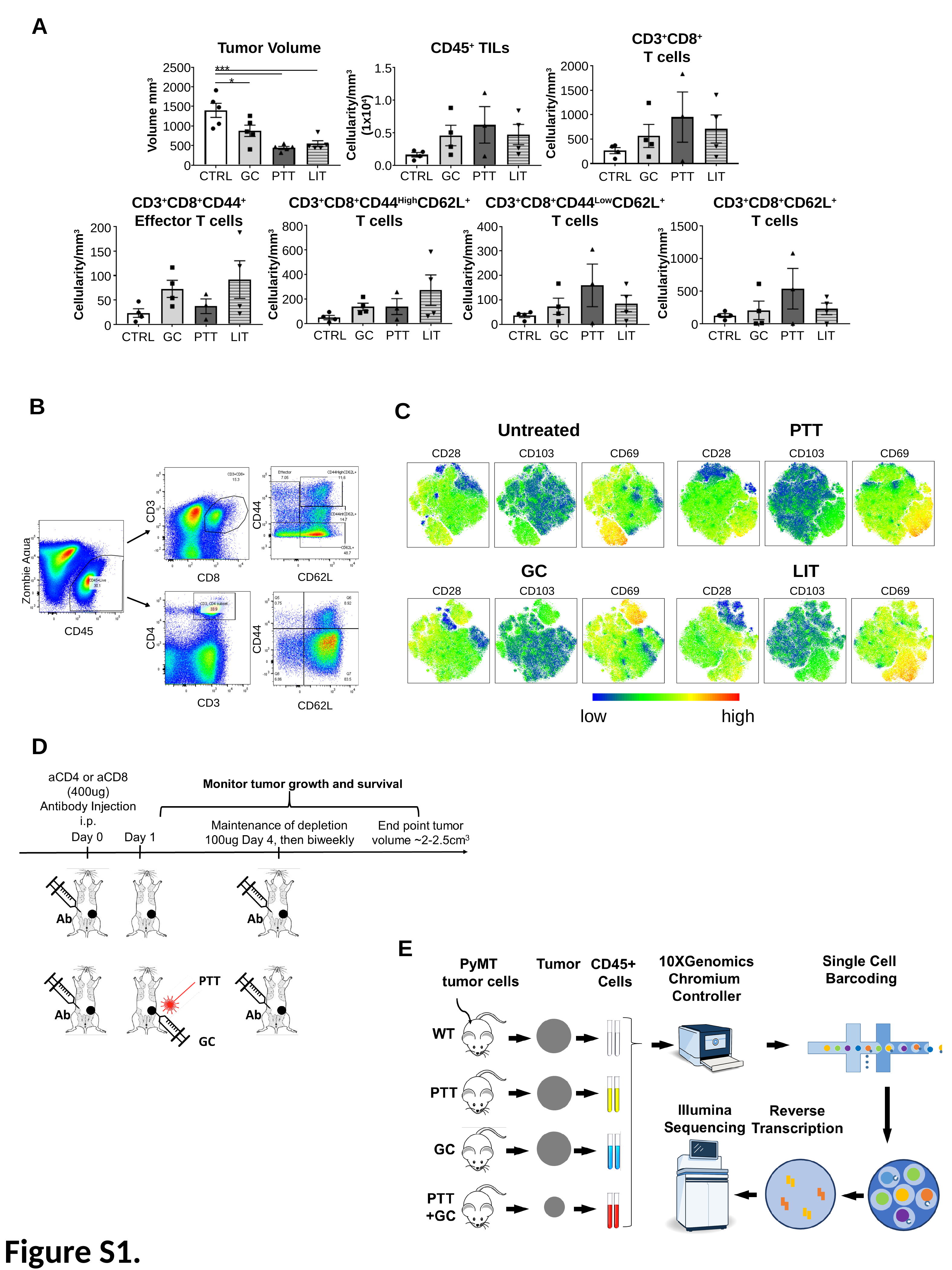

A
CD3^+^CD8^+^
T cells
Tumor Volume
CD45^+^ TILs
2000
2500
1.5
***
*
2000
1500
1.0
1500
Cellularity/mm^3^
(1x10^4^)
Cellularity/mm^3^
1000
Volume mm^3^
1000
0.5
500
500
0
0.0
0
CTRL
GC
PTT
LIT
CTRL
GC
PTT
LIT
CTRL
GC
PTT
LIT
CD3^+^CD8^+^CD44^+^
Effector T cells
CD3^+^CD8^+^CD44^High^CD62L^+^
T cells
CD3^+^CD8^+^CD44^Low^CD62L^+^
T cells
CD3^+^CD8^+^CD62L^+^
T cells
800
1500
400
200
600
300
150
1000
Cellularity/mm^3^
400
Cellularity/mm^3^
200
Cellularity/mm^3^
Cellularity/mm^3^
100
500
200
100
50
0
0
0
0
CTRL
GC
PTT
LIT
CTRL
GC
PTT
LIT
CTRL
GC
PTT
LIT
CTRL
GC
PTT
LIT
B
C
Untreated
PTT
CD28
CD103
CD69
CD28
CD103
CD69
GC
LIT
CD28
CD103
CD69
CD28
CD103
CD69
low
high
CD3
CD44
Zombie Aqua
CD62L
CD8
CD45
CD4
CD44
CD3
CD62L
D
E
Figure S1.

### Slide 2
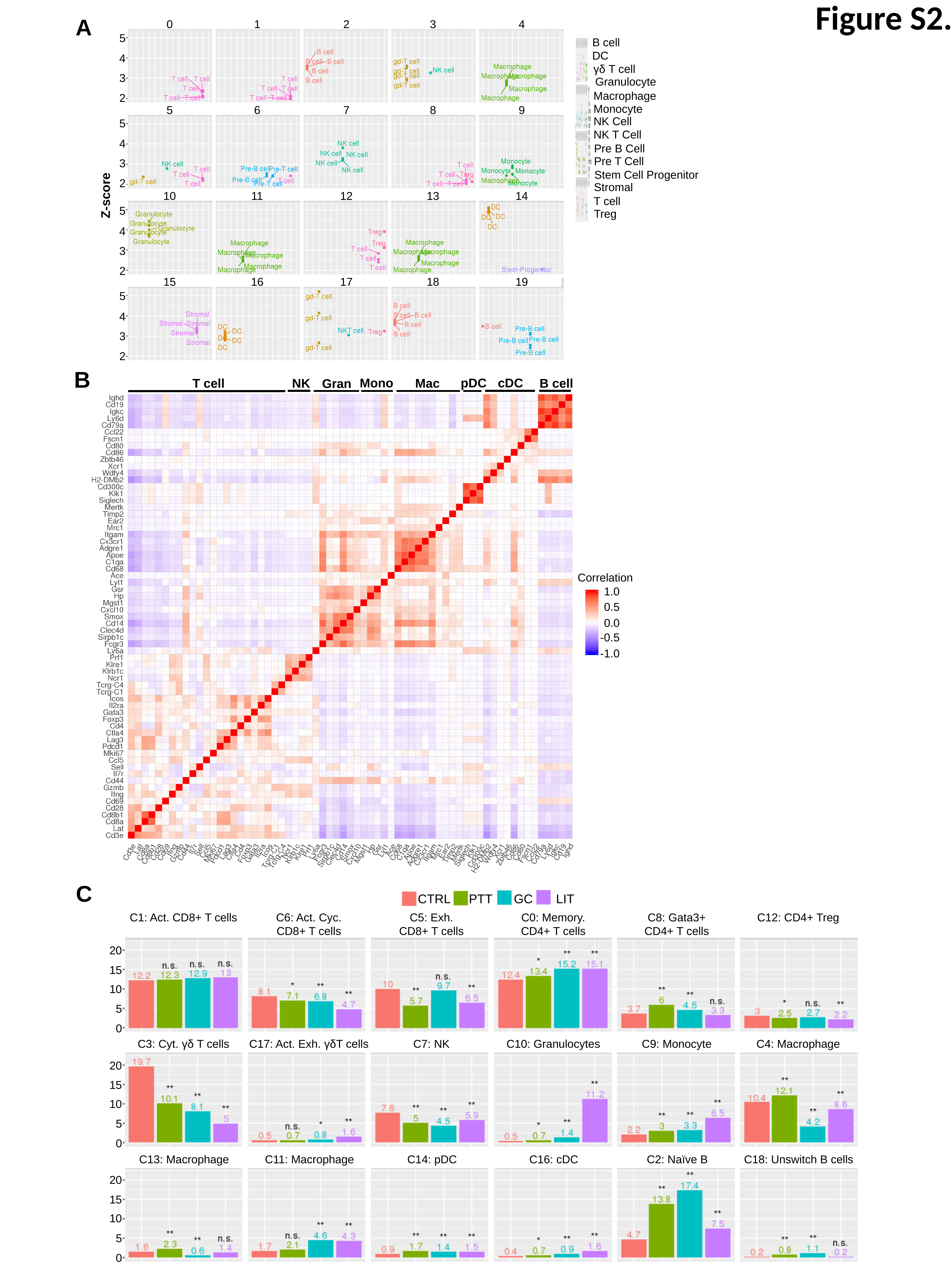

Figure S2.
A
0
1
2
3
4
5
4
3
2
B cell
DC
γδ T cell
Granulocyte
Macrophage
Monocyte
NK Cell
NK T Cell
Pre B Cell
Pre T Cell
Stem Cell Progenitor
Stromal
T cell
Treg
5
6
7
8
9
5
4
3
2
Z-score
10
11
12
13
14
5
4
3
2
15
16
17
18
19
5
4
3
2
B
pDC
cDC
Mono
Mac
B cell
NK
T cell
Gran
Correlation
1.0
0.5
0.0
-0.5
-1.0
C
CTRL
PTT
GC
LIT
C1: Act. CD8+ T cells
C6: Act. Cyc.
CD8+ T cells
C5: Exh.
CD8+ T cells
C0: Memory.
CD4+ T cells
C8: Gata3+
CD4+ T cells
C12: CD4+ Treg
20
15
10
5
0
C3: Cyt. γδ T cells
C17: Act. Exh. γδT cells
C7: NK
C10: Granulocytes
C9: Monocyte
C4: Macrophage
20
15
10
5
0
C13: Macrophage
C11: Macrophage
C14: pDC
C16: cDC
C2: Naïve B
C18: Unswitch B cells
20
15
10
5
0

### Slide 3
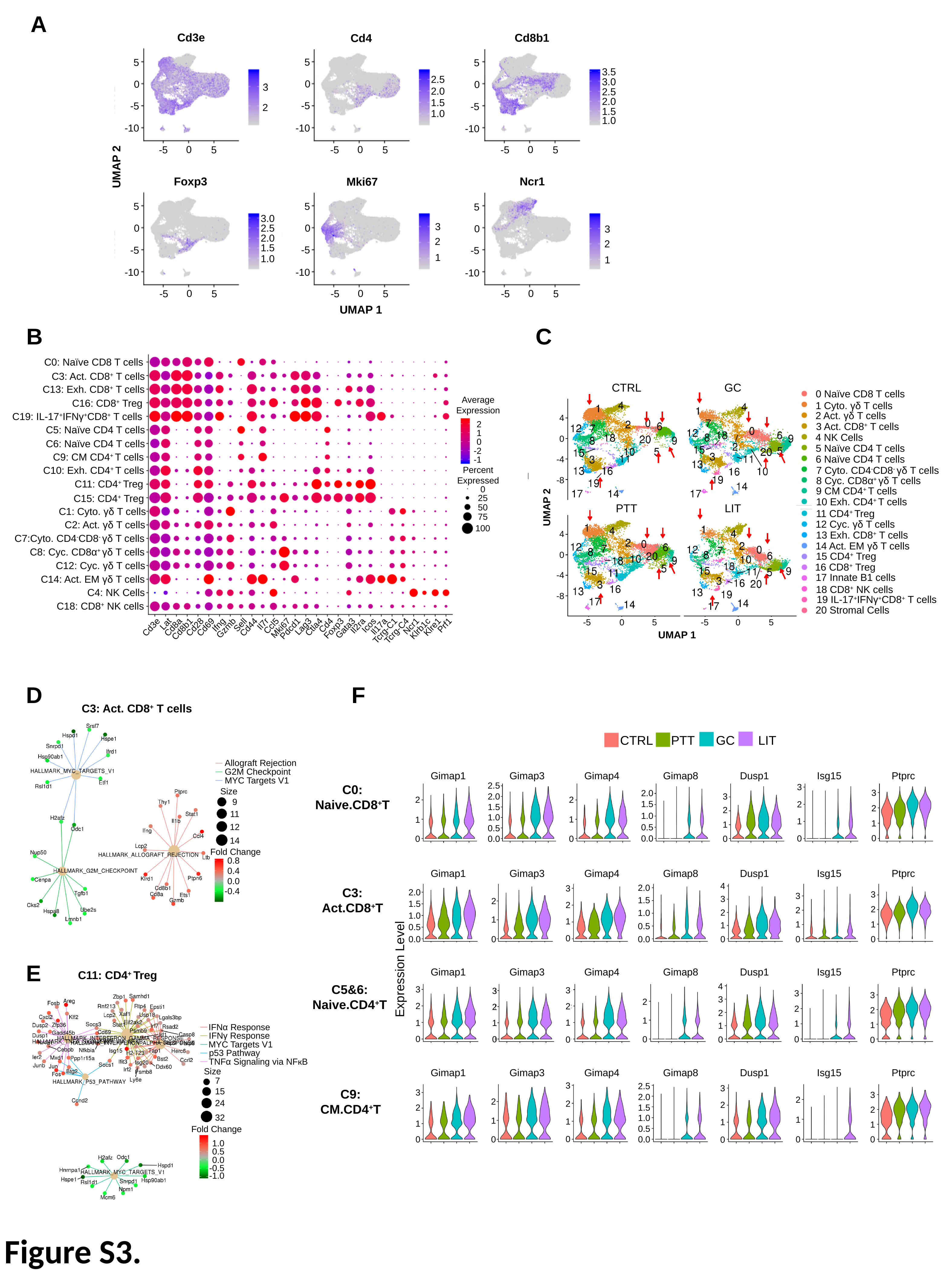

A
Cd3e
Cd8b1
Cd4
5
0
-5
-10
5
0
-5
-10
5
0
-5
-10
3.5
2.5
3.0
3
2
2.0
2.5
2.0
1.5
1.5
1.0
1.0
0
-5
5
0
-5
5
0
-5
5
UMAP 2
Foxp3
Ncr1
Mki67
5
0
-5
-10
5
0
-5
-10
5
0
-5
-10
3.0
3
2
1
2.5
3
2
1
2.0
1.5
1.0
0
-5
5
0
-5
5
0
-5
5
UMAP 1
B
C
C0: Naïve CD8 T cells
C3: Act. CD8^+^ T cells
C13: Exh. CD8^+^ T cells
Average
Expression
C16: CD8^+^ Treg
C19: IL-17^+^IFNγ^+^CD8^+^ T cells
2
C5: Naïve CD4 T cells
1
0
C6: Naïve CD4 T cells
-2
C9: CM CD4^+^ T cells
-1
C10: Exh. CD4^+^ T cells
Percent
Expressed
C11: CD4^+^ Treg
0
25
50
75
100
C15: CD4^+^ Treg
C1: Cyto. γδ T cells
C2: Act. γδ T cells
C7:Cyto. CD4^-^CD8^-^ γδ T cells
C8: Cyc. CD8α^+^ γδ T cells
C12: Cyc. γδ T cells
C14: Act. EM γδ T cells
C4: NK Cells
C18: CD8^+^ NK cells
Lat
Il7r
Ifng
Sell
Cd4
Icos
Prf1
Il2ra
Ccl5
Ncr1
Il17a
Lag3
Ctla4
Klre1
Cd3e
Cd8a
Cd28
Cd69
Cd44
Gzmb
Mki67
Gata3
Foxp3
Pdcd1
Klrb1c
Cd8b1
Tcrg-C1
Tcrg-C4
CTRL
GC
0 Naïve CD8 T cells
1 Cyto. γδ T cells
2 Act. γδ T cells
4
3 Act. CD8^+^ T cells
4 NK Cells
0
5 Naïve CD4 T cells
6 Naïve CD4 T cells
-4
7 Cyto. CD4^-^CD8^-^ γδ T cells
8 Cyc. CD8α^+^ γδ T cells
-8
9 CM CD4^+^ T cells
10 Exh. CD4^+^ T cells
PTT
LIT
UMAP 2
11 CD4^+^ Treg
12 Cyc. γδ T cells
4
13 Exh. CD8^+^ T cells
14 Act. EM γδ T cells
0
15 CD4^+^ Treg
16 CD8^+^ Treg
-4
17 Innate B1 cells
18 CD8^+^ NK cells
-8
19 IL-17^+^IFNγ^+^CD8^+^ T cells
20 Stromal Cells
-5
0
5
-5
0
5
UMAP 1
F
D
C3: Act. CD8^+^ T cells
Allograft Rejection
G2M Checkpoint
MYC Targets V1
Size
9
11
12
14
Fold Change
0.8
0.4
0.0
-0.4
CTRL
PTT
GC
LIT
Gimap1
Gimap3
Gimap4
Gimap8
Dusp1
Isg15
Ptprc
2.5
2.0
1.5
1.0
0.5
0.0
3
2
1
0
C0:
Naive.CD8^+^T
3
2
1
0
2.0
1.5
1.0
0.5
0.0
3
2
1
0
2
1
0
2
1
0
Gimap1
Gimap3
Gimap4
Gimap8
Dusp1
Isg15
Ptprc
4
3
2
1
0
2.0
1.5
1.0
0.5
0.0
3
2
1
0
C3:
Act.CD8^+^T
2.0
1.5
1.0
0.5
0.0
3
2
1
0
3
2
1
0
2
1
0
Expression Level
Gimap1
Gimap3
Gimap4
Gimap8
Dusp1
Isg15
Ptprc
4
3
2
1
0
C5&6:
Naive.CD4^+^T
3
3
2
1
0
3
2
1
0
3
2
1
0
3
2
1
0
2
1
0
2
1
0
Gimap1
Gimap3
Gimap4
Gimap8
Dusp1
Isg15
Ptprc
3
2
1
0
2.5
2.0
1.5
1.0
0.5
0.0
3
2
1
0
C9:
CM.CD4^+^T
3
2
1
0
3
2
1
0
2
1
0
2
1
0
E
C11: CD4^+^ Treg
IFNα Response
0
IFNγ Response
MYC Targets V1
p53 Pathway
TNFα Signaling via NFкB
Size
7
15
24
32
Fold Change
1.0
0.5
0.0
-0.5
-1.0
Figure S3.

### Slide 4
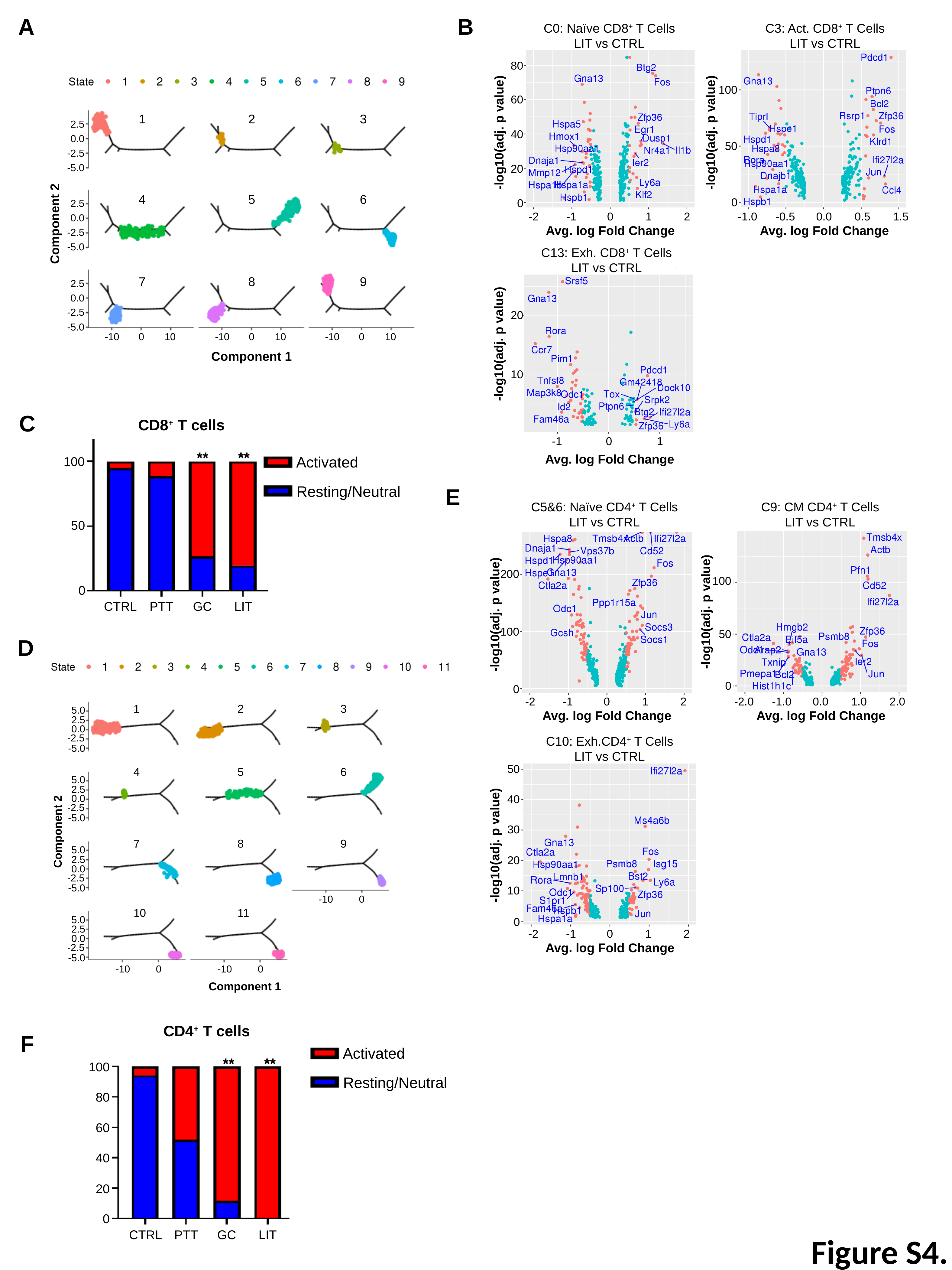

A
B
C0: Naïve CD8^+^ T Cells
LIT vs CTRL
C3: Act. CD8^+^ T Cells
LIT vs CTRL
80
100
60
-log10(adj. p value)
-log10(adj. p value)
40
50
20
0
0
-2
-1
0
1
2
-1.0
-0.5
0.0
0.5
1.5
Avg. log Fold Change
Avg. log Fold Change
C13: Exh. CD8^+^ T Cells
LIT vs CTRL
20
-log10(adj. p value)
10
-1
0
1
Avg. log Fold Change
1
3
5
7
9
State
2
4
6
8
1
2
3
2.5
0.0
-2.5
-5.0
4
5
6
2.5
0.0
-2.5
-5.0
Component 2
7
8
9
2.5
0.0
-2.5
-5.0
-10
0
10
-10
0
10
-10
0
10
Component 1
C
CD8^+^ T cells
**
**
Activated
Resting/Neutral
100
50
0
CTRL
PTT
GC
LIT
E
C5&6: Naïve CD4^+^ T Cells
LIT vs CTRL
C9: CM CD4^+^ T Cells
LIT vs CTRL
200
100
-log10(adj. p value)
-log10(adj. p value)
100
50
0
0
-2
-1
0
1
2
-2.0
-1.0
0.0
1.0
2.0
Avg. log Fold Change
Avg. log Fold Change
C10: Exh.CD4^+^ T Cells
LIT vs CTRL
50
40
30
-log10(adj. p value)
20
10
0
-2
-1
0
1
2
Avg. log Fold Change
1
4
7
10
2
5
8
11
State
D
3
6
9
1
2
3
5.0
2.5
0.0
-2.5
-5.0
4
5
6
5.0
2.5
0.0
-2.5
-5.0
Component 2
7
8
9
5.0
2.5
0.0
-2.5
-5.0
-10
0
10
11
5.0
2.5
0.0
-2.5
-5.0
-10
0
-10
0
Component 1
CD4^+^ T cells
Activated
Resting/Neutral
100
80
60
40
20
0
CTRL
PTT
GC
LIT
F
**
**
Figure S4.

### Slide 5
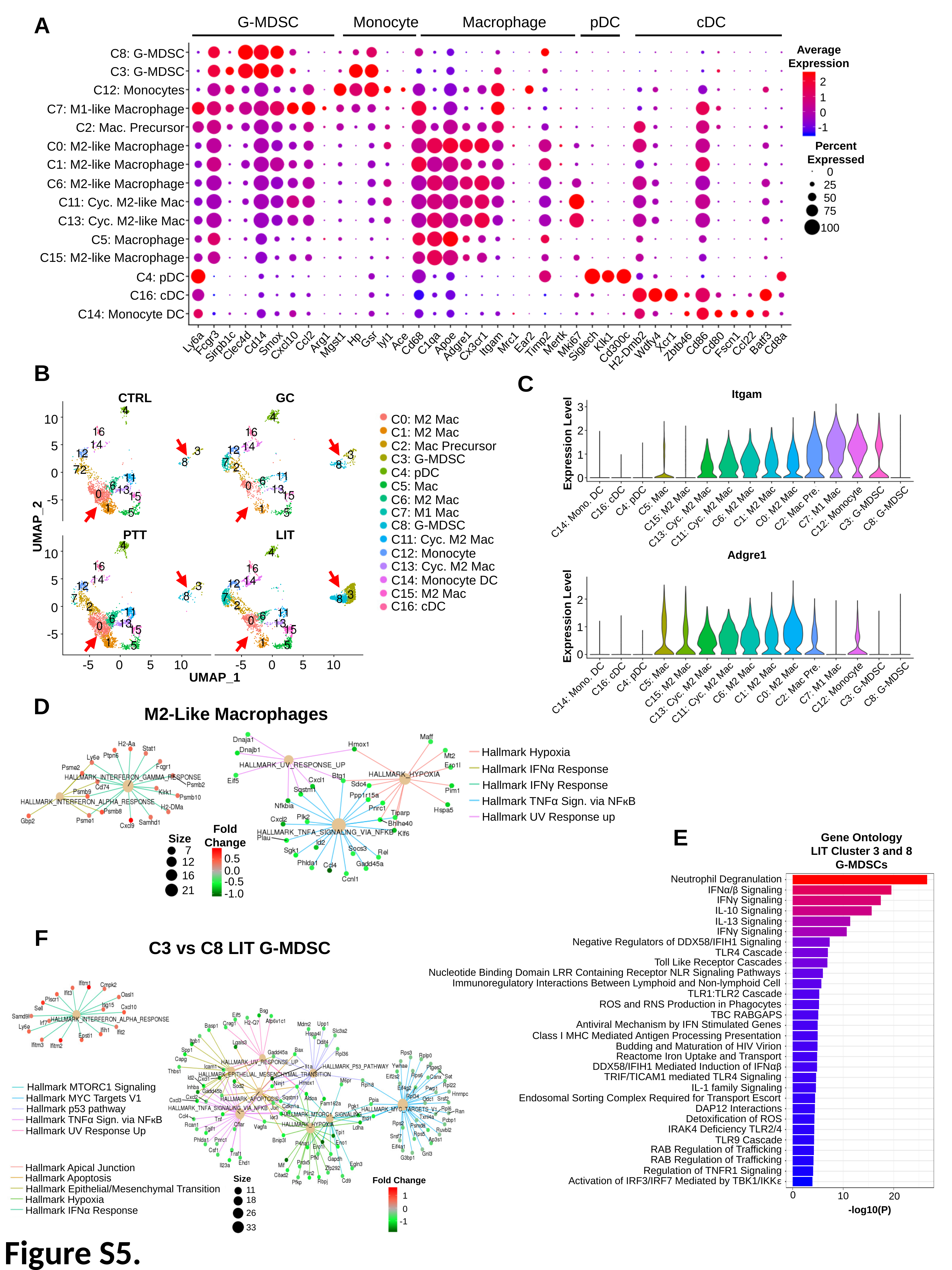

A
Macrophage
G-MDSC
pDC
Monocyte
cDC
Average
Expression
C8: G-MDSC
C3: G-MDSC
2
C12: Monocytes
1
C7: M1-like Macrophage
0
C2: Mac. Precursor
-1
C0: M2-like Macrophage
Percent
Expressed
C1: M2-like Macrophage
0
C6: M2-like Macrophage
25
50
C11: Cyc. M2-like Mac
75
C13: Cyc. M2-like Mac
100
C5: Macrophage
C15: M2-like Macrophage
C4: pDC
C16: cDC
C14: Monocyte DC
Hp
Gsr
lyl1
Ace
Ccl2
Klk1
Xcr1
Arg1
Ly6a
Ear2
Mrc1
Fcgr3
Mertk
Apoe
Cd14
Batf3
C1qa
Cd8a
Cd68
Cd86
Cd80
Smox
Ccl22
Itgam
Mki67
Timp2
Mgst1
Fscn1
Wdfy4
Cxcl10
Adgre1
Clec4d
Cx3cr1
Siglech
Zbtb46
Sirpb1c
Cd300c
H2-Dmb2
B
C
Itgam
3
2
Expression Level
1
0
C5: Mac
C4: pDC
C16: cDC
C6: M2 Mac
C1: M2 Mac
C0: M2 Mac
C7: M1 Mac
C2: Mac Pre.
C3: G-MDSC
C8: G-MDSC
C15: M2 Mac
C12: Monocyte
C14: Mono. DC
C11: Cyc. M2 Mac
C13: Cyc. M2 Mac
Adgre1
2
Expression Level
1
0
C5: Mac
C4: pDC
C16: cDC
C6: M2 Mac
C1: M2 Mac
C0: M2 Mac
C7: M1 Mac
C2: Mac Pre.
C3: G-MDSC
C8: G-MDSC
C15: M2 Mac
C12: Monocyte
C14: Mono. DC
C11: Cyc. M2 Mac
C13: Cyc. M2 Mac
CTRL
GC
C0: M2 Mac
C1: M2 Mac
C2: Mac Precursor
C3: G-MDSC
C4: pDC
C5: Mac
C6: M2 Mac
C7: M1 Mac
C8: G-MDSC
C11: Cyc. M2 Mac
C12: Monocyte
C13: Cyc. M2 Mac
C14: Monocyte DC
C15: M2 Mac
C16: cDC
10
5
0
-5
UMAP_2
PTT
LIT
10
5
0
-5
-5
0
5
10
-5
0
5
10
UMAP_1
D
M2-Like Macrophages
Hallmark Hypoxia
Hallmark IFNα Response
Hallmark IFNγ Response
Hallmark TNFα Sign. via NFкB
Hallmark UV Response up
Fold
Change
0.5
0.0
-0.5
-1.0
Size
7
12
16
21
E
Gene Ontology
LIT Cluster 3 and 8
G-MDSCs
Neutrophil Degranulation
IFNα/β Signaling
IFNγ Signaling
IL-10 Signaling
IL-13 Signaling
IFNγ Signaling
Negative Regulators of DDX58/IFIH1 Signaling
TLR4 Cascade
Toll Like Receptor Cascades
Nucleotide Binding Domain LRR Containing Receptor NLR Signaling Pathways
Immunoregulatory Interactions Between Lymphoid and Non-lymphoid Cell
TLR1:TLR2 Cascade
ROS and RNS Production in Phagocytes
TBC RABGAPS
Antiviral Mechanism by IFN Stimulated Genes
Class I MHC Mediated Antigen Processing Presentation
Budding and Maturation of HIV Virion
Reactome Iron Uptake and Transport
DDX58/IFIH1 Mediated Induction of IFNαβ
TRIF/TICAM1 mediated TLR4 Signaling
IL-1 family Signaling
Endosomal Sorting Complex Required for Transport Escort
DAP12 Interactions
Detoxification of ROS
IRAK4 Deficiency TLR2/4
TLR9 Cascade
RAB Regulation of Trafficking
RAB Regulation of Trafficking
Regulation of TNFR1 Signaling
Activation of IRF3/IRF7 Mediated by TBK1/IKKε
0
10
20
-log10(P)
F
C3 vs C8 LIT G-MDSC
Hallmark MTORC1 Signaling
Hallmark MYC Targets V1
Hallmark p53 pathway
Hallmark TNFα Sign. via NFкB
Hallmark UV Response Up
Hallmark Apical Junction
Hallmark Apoptosis
Hallmark Epithelial/Mesenchymal Transition
Hallmark Hypoxia
Hallmark IFNα Response
Size
11
18
26
33
Fold Change
1
0
-1
Figure S5.

### Slide 6
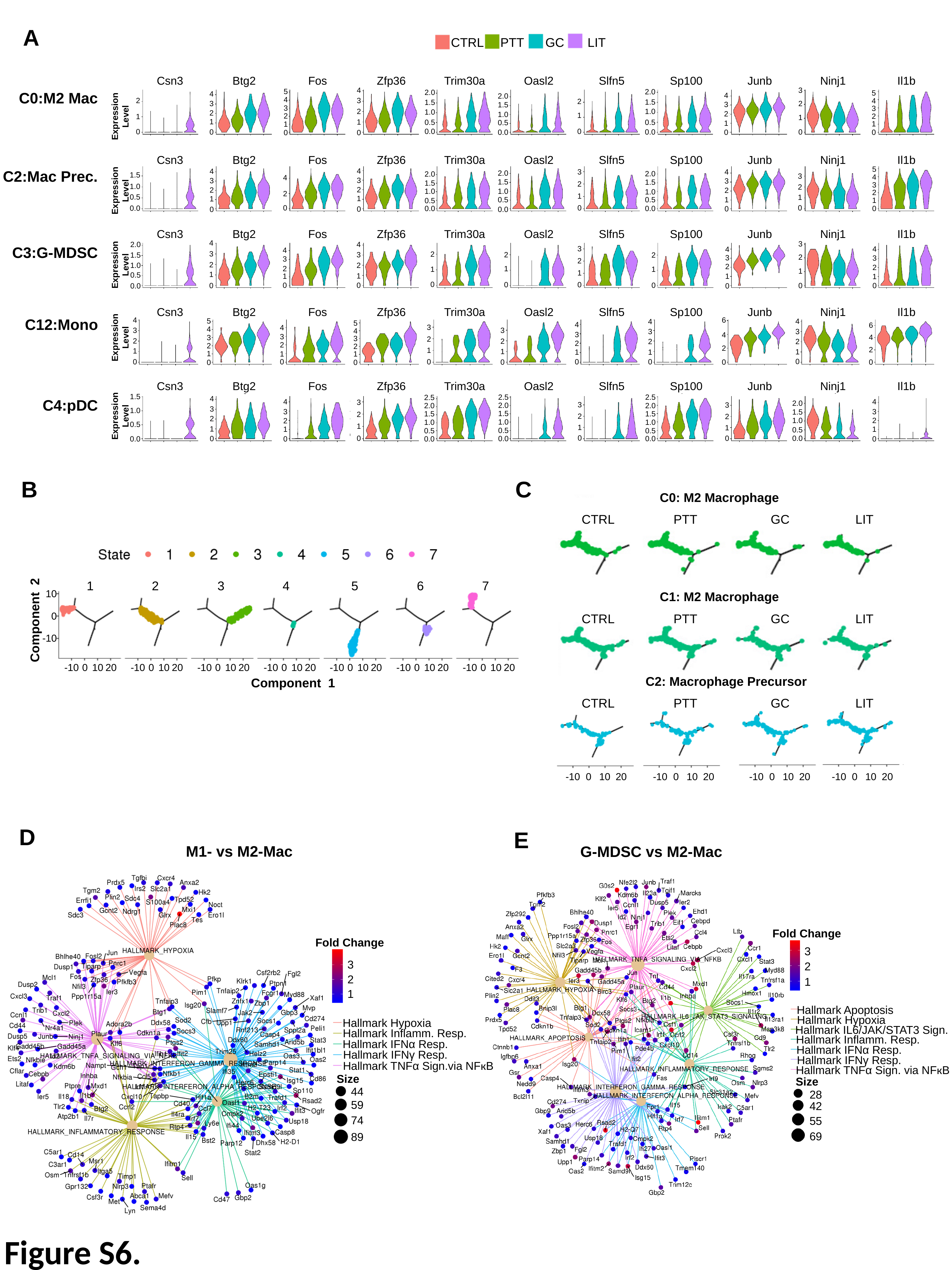

A
CTRL
PTT
GC
LIT
Csn3
Btg2
Fos
Zfp36
Trim30a
Oasl2
Slfn5
Sp100
Junb
Ninj1
Il1b
/
/
3
2
1
0
4
3
2
1
0
5
4
3
2
1
0
2.0
1.5
1.0
0.5
0.0
2.0
1.5
1.0
0.5
0.0
5
4
3
2
1
0
4
3
2
1
0
4
3
2
1
0
2.0
1.5
1.0
0.5
0.0
2
1
0
2
1
0
Expression
Level
Csn3
Btg2
Fos
Zfp36
Trim30a
Oasl2
Slfn5
Sp100
Junb
Ninj1
Il1b
/
/
/
5
4
3
2
1
0
4
3
2
1
0
4
3
2
1
0
3
2
1
0
4
3
2
1
0
2.0
1.5
1.0
0.5
0.0
2.0
1.5
1.0
0.5
0.0
2.0
1.5
1.0
0.5
0.0
1.5
1.0
0.5
0.0
3
2
1
0
4
2
0
Expression
Level
Csn3
Btg2
Fos
Zfp36
Trim30a
Oasl2
Slfn5
Sp100
Junb
Ninj1
Il1b
/
/
/
/
4
3
2
1
0
3
2
1
0
4
3
2
1
0
2.0
1.5
1.0
0.5
0.0
3
2
1
0
4
3
2
1
0
4
3
2
1
0
4
3
2
1
0
2
1
0
2
1
0
2
1
0
Expression
Level
Csn3
Btg2
Fos
Zfp36
Trim30a
Oasl2
Slfn5
Sp100
Junb
Ninj1
Il1b
/
4
3
2
1
0
6
4
2
0
5
4
3
2
1
0
4
3
2
1
0
4
3
2
1
0
5
4
3
2
1
0
4
3
2
1
0
6
4
2
0
3
2
1
0
4
3
2
1
0
3
2
1
0
Expression
Level
Csn3
Btg2
Fos
Zfp36
Trim30a
Oasl2
Slfn5
Sp100
Junb
Ninj1
Il1b
1.5
1.0
0.5
0.0
2.5
2.0
1.5
1.0
0.5
0.0
2.5
2.0
1.5
1.0
0.5
0.0
2.5
2.0
1.5
1.0
0.5
0.0
4
3
2
1
0
4
3
2
1
0
4
3
2
1
0
3
2
1
0
2.0
1.5
1.0
0.5
0.0
3
2
1
0
3
2
1
0
Expression
Level
C0:M2 Mac
C2:Mac Prec.
C3:G-MDSC
C12:Mono
C4:pDC
B
C
C0: M2 Macrophage
CTRL
PTT
GC
LIT
C1: M2 Macrophage
CTRL
PTT
GC
LIT
C2: Macrophage Precursor
CTRL
PTT
GC
LIT
-10
0
10
20
-10
0
10
20
-10
0
10
20
-10
0
10
20
1
2
3
4
5
6
7
10
0
Component 2
-10
-10
0
10
20
-10
0
10
20
-10
0
10
20
-10
0
10
20
-10
0
10
20
-10
0
10
20
-10
0
10
20
Component 1
D
E
G-MDSC vs M2-Mac
Fold Change
3
2
1
Hallmark Apoptosis
Hallmark Hypoxia
Hallmark IL6/JAK/STAT3 Sign.
Hallmark Inflamm. Resp.
Hallmark IFNα Resp.
Hallmark IFNγ Resp.
Hallmark TNFα Sign. via NFкB
Size
28
42
55
69
M1- vs M2-Mac
Fold Change
3
2
1
Hallmark Hypoxia
Hallmark Inflamm. Resp.
Hallmark IFNα Resp.
Hallmark IFNγ Resp.
Hallmark TNFα Sign.via NFкB
Size
44
59
74
89
Figure S6.

### Slide 7
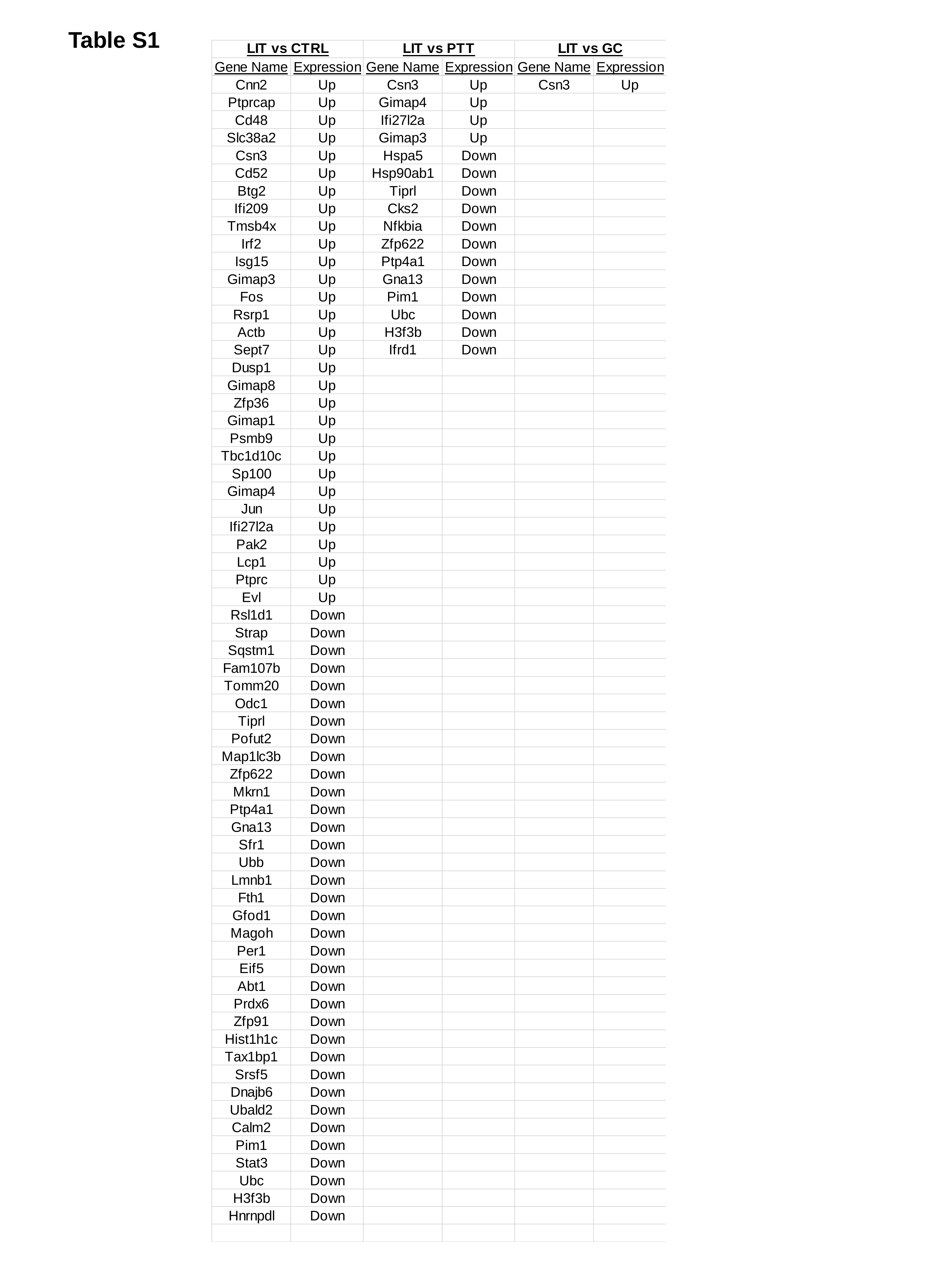

Table S1

### Slide 8
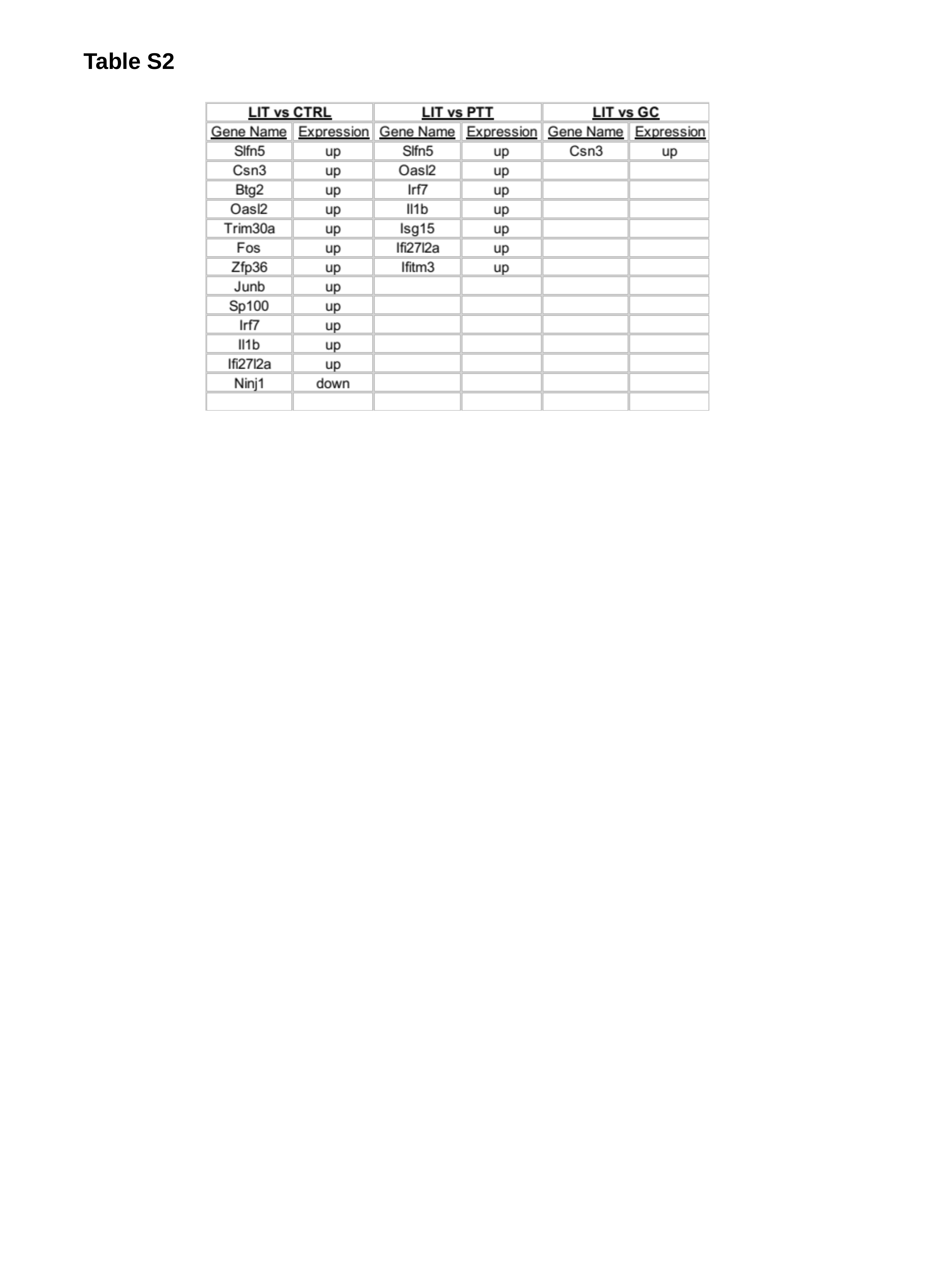

Table S2
